# Antarctic biosecurity policy effectively manages the rates of alien introductions

**DOI:** 10.1101/2024.10.26.620391

**Authors:** Rachel I. Leihy, Melodie A. McGeoch, David A. Clarke, Lou Peake, Yehezkel Buba, Jonathan Belmaker, Steven L. Chown

**Affiliations:** Securing Antarctica’s Environmental Future, School of Biological Sciences, Monash University, Victoria, 3800, Australia; Arthur Rylah Institute for Environmental Research, Department of Energy, Environment, and Climate Action, Heidelberg, Victoria, 3084, Australia; Securing Antarctica’s Environmental Future, Department of Environment and Genetics, La Trobe University, Victoria, 3086, Australia; School of Zoology, Faculty of Life Sciences, Tel Aviv University, Tel Aviv, Israel; Steinhardt Museum of Natural History, Tel Aviv University, Tel Aviv, Israel

## Abstract

Reducing the rates and impacts of biological invasions is a major policy goal of international biodiversity agreements. Yet the extent to which this goal is being achieved and the agreements hence successful in this respect remains unclear. Here we use a comprehensive record of alien species introduction in the terrestrial Antarctic, including its surrounding Southern Ocean Islands, spanning 115 years (1900–2015), to quantify the impact of biosecurity policy on alien species introduction rates in the region, where invasive alien species are a primary environmental conservation threat and management priority. We show that although many parts of the Antarctic have been colonised by non-indigenous taxa, recent rates of introduction appear to be slowing or static in most parts, compared with increases in the past. Our results vindicate the regional Antarctic focus on biosecurity measures, but also demonstrate the need for stricter enforcement due to rapid socio-environmental changes.

**Three key points:**

- Biological invasions present a large and growing threat to Antarctic ecosystems under climate change and expanding human activity
- Over the 20^th^ century, there was no trend in alien species introduction rate in five Antarctic regions and a significantly increasing trend in the remaining five regions
- Despite this variation, in most regions, the number of introductions remains low, indicating that Antarctic biosecurity has been effective at slowing the rate of introduction

## Introduction

Antarctica and the surrounding Southern Ocean Islands (SOIs) have a relatively recent history of human occupation (the continent itself was only discovered ∼200 years ago), and far less human activity than is typical of most continents and islands elsewhere (Chown et al., 2017). Yet the region is not free of alien species and their impacts. Beginning with sealers and whalers, and followed by explorers, scientists and tourists, human visitors have introduced many species, including microbes, plants, invertebrates and vertebrates (Frenot et al., 2005; Headland, 2012; Hughes et al., 2015). Several have had extensive ecological impacts, including population extirpation (Connan et al., 2022; Lebouvier et al., 2011), and community-wide impacts on ecosystem structure and functioning (Bartlett et al., 2023; Chown et al., 2022). Indeed, ∼ 13% of established species in the region are known to be invasive (Leihy et al., 2023) (see Table S1 for glossary). Moreover, the numbers of species introduced across the region (now totalling over 1200) are closely linked to local climates and numbers of visitors to a given location (Leihy et al., 2018, 2023). Several studies have also demonstrated, or strongly implied, increasing population level impacts of these species with changing climates (Barbraud et al., 2021; McClelland et al., 2018). Hence, there is concern about the growing likelihood of establishment success and impact as climates change in the Antarctic (Bergstrom 2022; Convey & Lebouvier 2009; Hughes et al., 2020). This mirrors the significant concern raised by the recent, first global assessment of invasive alien species, demonstrating the mounting threat that invasive species pose to biodiversity and ecosystems (Roy et al., 2024).

Both globally (Butchart et al., 2010; CBD, 2022), and in the Antarctic (Antarctic Treaty Secretariat, 1991; CEP, 2019), policymakers have established policies to reduce the rates and impacts of biological invasions and goals to measure the success of these policies. In the Antarctic region, biosecurity policy provisions have a long history (∼ 60 years). From the 1990s onwards, however, measures became much stricter. For the Antarctic continent, the Protocol on Environmental Protection to the Antarctic Treaty (Antarctic Treaty Secretariat, 1991) (hereafter, the ‘Protocol’) and its Annex II effectively prohibited the introduction of non-native species, with the Committee for Environmental Protection (CEP) making alien species impacts a growing priority from the mid-2000s onwards (Rogan-Finnemore, 2008). Across the SOIs, governed by a variety of states (De Villiers et al., 2006), specific biosecurity legislation began to be applied at about the same time (Table S2), although new alien species continued to establish across the region (Greve et al., 2017; Hughes et al., 2020; Lebouvier et al., 2011).

Despite growing policy attention and urgent calls for further action given new alien species detections over the past ∼20 years (McGeoch et al., 2015), few attempts have been made to assess the efficacy of Antarctic biosecurity provisions. Given that alien species are the top priority of Antarctica’s CEP (CEP, 2023), and among the most significant concerns for conservation of the SOIs, and that rates of arrival of alien species are widely thought to be likely to increase in response to changing climates and growing visitor numbers (Hughes et al. 2020, 2021), this situation is surprising. To some extent, it reflects the general difficulty of assessing invasive species policy efficacy (Vicente et al., 2022). It may also reflect growing recognition that simply using the observed number of new introductions is not adequate for assessing trends in alien species introductions (McGeoch et al., 2023). Specifically, lags in alien species detection, varying investment in surveillance efforts over time and between regions, and sparse data on species impacts mean that producing robust estimates of rates of species invasion is challenging (Buba et al., 2024; Solow & Costello, 2004). In consequence, both for the Antarctic region and elsewhere, the efficacy of policy on altering the rates of alien species introductions remains largely unknown (or problematic if the biases have not been adequately considered).

Here, using the most contemporary and spatially-explicit compilation of alien species data available for the region (Leihy et al., 2023), and newly-developed, appropriate methods to assess rate variation, we investigate the extent to which policy interventions may have affected alien species introduction rates across the Antarctic between 1900 and 2015. We expect alien introduction rates to have increased through time, as they have globally (Bonnamour et al., 2021; Roy et al., 2024), and that they will continue to do so because of changing climates and growing human visitor numbers to the region from scientific and tourism interest (Duffy et al., 2017; Hughes et al., 2020, 2021).

## Methods

### Introduced species data

An open dataset on the identity, localities, and dates of introduction of introduced and invasive alien species was recently published for Antarctica and the SOIs (Leihy et al., 2023). This dataset includes records of species or other taxonomic levels (hereafter, simply referred to as species) introduced to Antarctic places outside their native range, including vagrants, domestic species, established and naturalised aliens, invasive species, and species that were introduced to the region but have since died out or been eradicated. It includes 3066 records of terrestrial and freshwater aliens, primarily vertebrates (mammals, birds, freshwater fish), invertebrates (insects, spiders, annelids, springtails, molluscs), and vascular plants. Other groups, such as fungi, alien marine species and microorganisms, were not included in the introduced and invasive alien species dataset (Cowan et al., 2011; Leihy et al., 2023; McCarthy et al., 2019).

The specificity of the locality information in the alien species dataset varies across records, with species reported at an individual island-, archipelago-, or subregional-scale (e.g., ‘Possession Island’ vs. ‘Crozet archipelago’). To reduce spatial bias in the data, we pooled data within each Southern Ocean archipelago, and for the Antarctic continent (areas south of 60°S) (hereafter, subregions), and retained only the first observation record per species and subregion (see Table S3 for subregional summary information). This standardises the spatial and taxonomic resolution of these data, but reduces the total number of introductions where species may have been introduced multiple times to sites within the same subregion. We excluded data from Islas Malvinas/Falkland Islands because more than half of the records have incomplete introduction dates. For other subregions, we added to the existing dataset (Leihy et al., 2023) through a search of the scientific and grey literature for missing dates of introduction to improve the completeness of the records. In total, we added 6 new alien species records and 108 introduction dates to existing records (Table S4).

The alien species dataset includes information on the date that species were first observed (i.e. discovered) at Antarctic localities and the year of the first published record on the discovery (Table S1). In the pooled dataset, the first observation and publication dates were selected per species and subregion. First published records were always recorded as a year, however, ranges of years were sometimes reported for first observation dates (e.g., ‘1950– 1953’; relevant to 10.2% of records). We followed guidelines provided by Seebens et al. (2017) and Dyer et al. (2017) to address these date ranges by randomly selecting a year within the range to avoid arbitrary peaks at the beginning or middle of the range.

While species have been introduced to the sub-Antarctic from as early as the 13th century (Russell et al., 2020), we selected 1900 as a uniform start date to compare trends across subregions with different histories of human occupation. In total, we included data for 1180 alien species populations, and excluded 155 populations introduced before 1900, 7 records from after 2015, and 277 populations with no introduction dates available (Table S3). We pooled the number of alien species observations per five year period between 1900 and 2015 to reduce the influence of inter-annual variation in survey effort due to the remoteness of these regions and access limitations. Five-year periods with no new alien species observations were assigned a zero value for analysis. We excluded recent observations from 2016 onwards to reduce the publication lag effect caused by a delay between the first observation of a newly discovered alien and the first published record (on average, 7.23 years; Figure 1).

**Figure 1.**
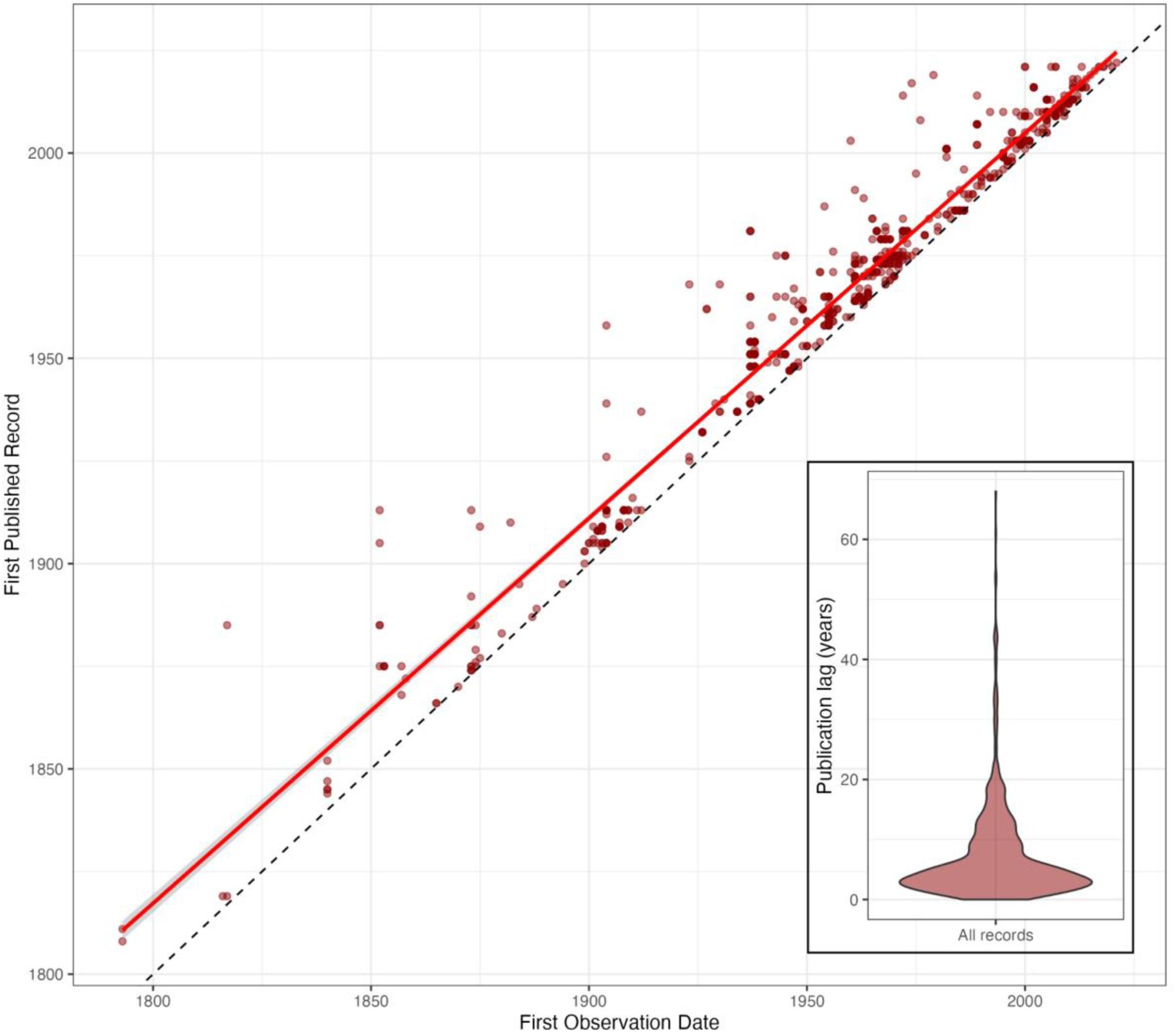
Publication lag reporting Antarctic alien species discoveries. Time lag between alien species discovery (first observation date) and first publication date in the scientific or grey literature for Antarctic and Southern Ocean Island alien species records between 1793 and 2022. The linear relationship between dates (solid red line; f*(x)* = 1.02*x* – 46.25; r^2^ = 0.96, df = 866; 95% confidence intervals in shaded grey area) was used to estimate first observation dates for records with publication dates only. Dashed line indicates the line of parity. Inset shows the distribution of records based on their publication lag in years (i.e., difference between first published and first observed record dates). On average, publications of records occurred 7.23 years after first discovery in the Antarctic region (min = 0 years, max = 68 years). For 15.4% of records with a range of dates recorded for the first observation date (e.g., “1950-1951”), we randomly selected a year within the range, following guidelines (Dyer et al., 2017; Seebens et al., 2017).

The completeness of the alien species data will influence the accuracy of the estimated introduction rate trends for Antarctica. Dataset completeness may be influenced by: missing introduction date and survey effort information, under-reporting on alien species due to limited survey effort or, in some cases, their perceived social or economic value at the time of introduction (such as ornamental and agricultural plant species, and pets), time lags between on-ground activities and subsequent publications, language barriers to data collection, and missed publications that are not widely cross-referenced (Leihy et al., 2023). Taken together, these factors may cause the underestimation of alien species introduction rates, especially in earlier periods of assessment. These sources of bias are common to most alien species analyses (McGeoch et al., 2023; Pagad et al., 2018), but may be particularly important in an Antarctic context where visitation to these remote places has been sporadic due to their inaccessibility (e.g., Heard Island), and due to the relatively low overall number of alien and indigenous species (i.e., any new species may be a significant discovery).

To estimate a first observation date for records with missing first observation dates, we modelled the linear relationship between first observation and first published record dates for records with both dates available (Figure 1). This model was used to estimate first observation dates for records with only a first publication date (relevant to 11.4% records).

### Survey effort proxy data

Survey effort affects detected rates of introduction (Buba et al., 2024; Wonham & Pachepsky, 2006), however, standard approaches to measuring and reporting survey effort for biological introductions have yet to be developed (McGeoch et al., 2023). Here, we use the number of scientific publications with a field research component per subregion and 5-year period as a proxy for survey effort. This proxy assumes that the number of publications based on field research is related to the extent of environmental observation at sites and the probability of observing a new introduction. We assume, from extensive experience in the region by one of the authors (Chown & Language, 1994), that researchers would report new “unusual” species during the collection of general environmental data, regardless of the scientific discipline.

Publication data were collected from a systematic Web of Science search with Boolean strings using combinations of regional place names (Table S5). Publications based on remote-sensed data, literature reviews, and meta-analyses without a field component were excluded manually for each subregion, excluding Antarctica. The Web of Science search for Antarctic publications returned more than 30,000 publications for the continental subregion alone (Table S5). To estimate the number of relevant publications for Antarctica, we used the temporal trend in the number of scientific publications with a field research component versus the total number of publications in the Web of Science search (e.g., our retention rate) for SOI subregions processed manually to predict the Antarctic proxy for survey effort. The relative number of SOI publications based on field research declined between 1900 and 2023 due to the increasing prevalence of remote-sensed data, literature reviews, and meta-analyses without a field component (*f(x)*= −0.001 *year* + 2.47).

### Introduction rate models

Naïve model: First, to examine trends in the rate of alien species introductions over the last century, we fit Poisson Generalised Linear Models (GLMs) to the number of new detected alien species per five-year period and the time since 1900. This naïve introduction rate model assumes perfect detection of new alien species over time. Trends were fitted independently for each subregion, then p-values were adjusted for multiple comparisons using a False Discovery Rate correction to reduce Type 1 errors. We also fit Poisson GLMs to the survey effort trends over time, to compare the naïve introduction rate models to survey effort trends in each subregion. These trends can be qualitatively compared and interpreted to estimate the risk of underestimating the rate of new alien species observations over time (McGeoch et al., 2015).

Solow and Costello model: Second, we fit the Solow and Costello (2004) model to the introduction rate data, using the alien package in R (v. 1.0.1; Buba, 2024). These models estimate the rate of introduction of alien species from the discovery record, allowing for a monotonic increase in mean introduction rate over time (Solow & Costello, 2004). These models assume that the overall detection probability increases exponentially over time, and that each species detection also increases following its introduction due to growth in population abundance of the species and the detection likelihood (see more detail about the growth parameter, γ_2_, below) (Buba et al., 2024).

Sampling model: Third, we used a sampling model approach to quantitatively incorporate information about temporal trends in survey effort into the introduction rate model (Buba et al., 2024). Previous studies have shown that the naïve modelling approach typically overestimates introduction rates, particularly for shorter time series (< 50 years) or where there has been imperfect and variable sampling effort over time (Buba et al., 2024). Likewise, the Solow and Costello (2004) models assume the temporal trend in sampling effort is monotonic, yet in practice, sampling effort may vary over time (Buba et al., 2024). Human activity across the broader Antarctic region has varied markedly over the last century; characterised by sporadic scientific expeditions and commercial whaling and sealing activities in the first half of the century (Chown et al., 2005), before routine scientific and tourism activities commenced after World War II (Table S2). We, therefore, expect that there has been variable survey effort over time in these regions. Here, we use the number of publications based on field research per 5-year period as an independent proxy for survey effort (described above) in the sampling model. The survey effort proxy was standardised using the scale function in R. Sampling models were run using the snc function in the alien R package (v. 1.0.2.9; Buba, 2024). For each subregion, we fit the sampling model using time (*t*; number of years from the start of the timeseries) as a predictor of the 5-year introduction rate (μ_t_), and the rescaled survey effort proxy as a predictor of the 5-year probability of detection (Π_st_). For each region, we fit models using both the linear and exponential functions for the introduction rate formula, and used AIC values, derived from the maximum log-likelihoods, to select the best-fitting model (Table S6). Likewise, we fit models with and without the growth parameter (γ_2_) for the probability of detection (Π_st_) model, and used AIC to select the best-fitting model (Table S6). This parameter (also in the Solow and Costello model) specifies the detection model so that the detection probability increases over time due to growth in population abundance of the species and the detection likelihood. Removing the growth parameter simplifies the model, so that detection probability only depends on variation in the survey effort proxy over time. The Models that failed to converge when the maximum number of iterations was reached (10,000) were excluded (i.e., Heard and McDonald Islands and South Georgia and the South Sandwich Islands).

All introduction rate models (naïve, Solow and Costello, and sampling) were evaluated using bias and mean squared error (MSE) metrics. All analyses were performed in R (v.4.2.2; R Core Team, 2022), using the alien (Buba, 2024), rjags (v. 4-15; Plummer, 2023), ggplot2 (v.3.4.0; Wickham, 2016) and gridExtra (v.2.3; Auguie, 2017) packages.

## Results

Three modelling approaches were used to estimate the rates of alien species introduction: i) a simple ‘naïve’ model that disregards sampling effects, ii) a previous model that assumes that the probability of detecting introduced species increases monotonically and exponentially over time since introduction (Solow & Costello, 2004), and iii) a newly developed sampling model that incorporates independent data on survey effort over time to define the detection probability (Buba et al., 2024). The performance of these models differs under different local contexts (Buba et al., 2024), therefore, using and comparing the outputs of multiple models as we do here provides a robust estimation of introduction trends across the region.

### Naïve model assuming perfect detection

Rates of alien species introductions have varied across the Antarctic region. Using the naïve model, introduction rates increased significantly in five subregions: Antarctica (the continent and islands immediately surrounding it), Crozet Islands, Amsterdam and St. Paul Islands, Prince Edward Islands, and the Tristan da Cunha group (Figure 2; Table S7). For the remaining five Antarctic subregions (Heard and McDonald Islands, Kerguelen Islands, Macquarie Island, New Zealand sub-Antarctic Islands, and South Georgia and the South Sandwich Islands), there was no trend in introduction rate over time (Figure 2; Table S7). Over the same period (1900-2015), survey effort (measured using a scientific publication proxy, see Methods) increased significantly in all Antarctic subregions (Figure 2).

**Figure 2.**
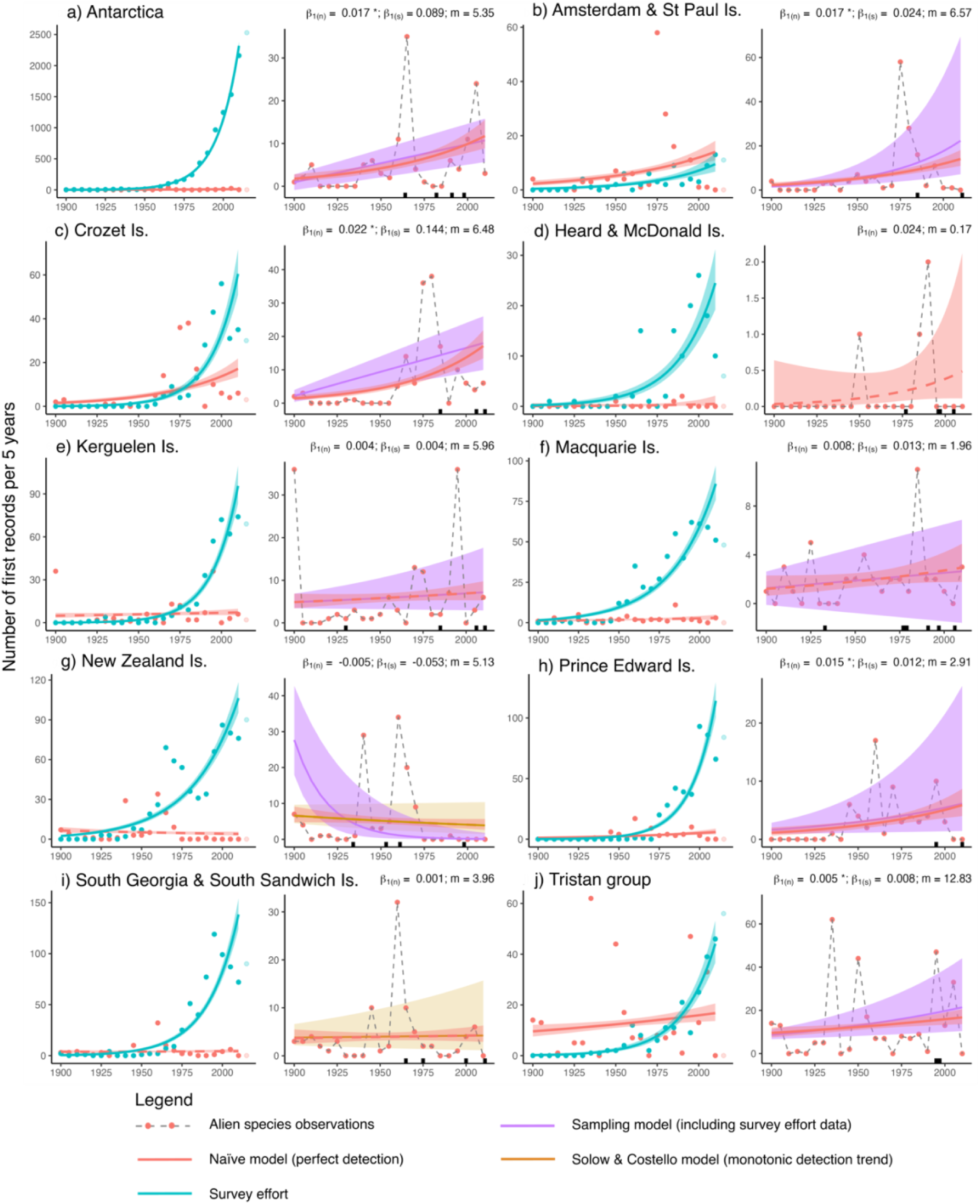
Rates of Antarctic alien introductions. Trends in alien species introductions and in survey effort across the broader Antarctic region, estimated using three models. The first panel of each subregion shows the naïve model (red line) of the number of alien species observations per five-year period (red points), assuming perfect detection of new aliens, and the alien survey effort trend (blue line) based on the number of publications with a field research component (blue points), fitted with Poisson Generalised Linear Models. Records from 2015 onwards (transparent points) were not included in models due to a publication lag between new observations and published reports. Non-significant trends after False Discovery Rate correction indicated with a dashed line (d,e,f,g,i). In the second panel for each subregion, trends show the naïve model (red line), sampling model (purple line) incorporating survey effort data, and Solow and Costello (SC) model (yellow line). The SC model is produced here only for the New Zealand sub-Antarctic islands (g) and South Georgia and South Sandwich islands (i) to improve visualisation (due to their poor fit with the observed data across most regions, see Table S8 and Figure S1). In all plots, shading indicates 95% confidence intervals. Values indicate the rate of change in number of alien species per five-year period for the naïve model (β_1(n)_) and sampling model (β_1(s)_), and mean number of alien species estimated from the naïve model, averaged across the time-series (*m*). Black x-axis bars indicate when key conservation policies were adopted (Table S2).

The estimated mean number of alien species introductions across the Antarctic region typically ranged between 2 – 7 new alien species per five-year period (Table S8). For the Heard and McDonald Islands, however, the average introduction rate was lower (∼ 0.17 new aliens per five year period), whereas the naïve model predicted fastest introduction rates in the Tristan da Cunha group compared to other subregions (mean 12.83 new aliens per five year period; Table S8).

### Solow and Costello model assuming increasing survey effort

For most Antarctic subregions, the Solow and Costello model (which assumes that survey effort increases monotonically with time) were a poor fit to the observed introduction rate data and produced highly-biased estimates of introduction rate compared to the naïve and sampling models (Figure 2; Table S8). For South Georgia and the South Sandwich Islands and the New Zealand sub-Antarctic islands, however, the Solow and Costello and naïve model results were similar, estimating a constant introduction rate of ∼ 4 and ∼ 5 new species per five year period, respectively (Figure 2g, i).

### Sampling model incorporating survey effort data

The sampling model, which incorporates survey effort data, closely resembles the naïve model for most Antarctic subregions (Figure 2), with similar error and bias (Table S8). The rate of change in number of alien species per five year period varied across subregions, between - 0.028 for the New Zealand sub-Antarctic islands and 0.144 for Crozet (Figure 2). This agreement between the sampling and naïve model provides support for the naïve model, and, importantly, suggests that detection probabilities have been generally high across the region (an assumption of the naïve model).

In most Antarctic subregions, peaks in observed alien species introductions occurred in the middle of the 20th century, exceeding naïve and sampling model expectations (Figure 2). Exceptions to these general patterns were the Heard and McDonald Islands, where observed and simulated first record rates were low over time (< 1 new alien per five year period) (Figure 2d), the New Zealand sub-Antarctic islands, where there has been a slight declining observed introduction trend (Figure 2g), and the Kerguelen Islands which had an early peak in alien species discoveries (following the outcomes of the Gauss Expedition 1901–1903) (Figure 1e).

Among the five subregions with increasing introduction trends, lower than expected numbers of observed alien species were detected from 2000 onwards in three regions: Amsterdam and St. Paul, Crozet, and the Prince Edward Islands (Figure 2b,c,h). This recent downturn suggests that the increase in human activity in recent decades, alongside an increase in survey effort, has not resulted in an increase in the number of new species discoveries in these regions. For Tristan da Cunha and Antarctica, however, introduction rates have increased recently, in some years exceeding expectations (Figure 2a, j).

## Discussion

Across the Antarctic region, rates of alien species introductions have remained static or increased slightly over the 20th century at between −0.028 and 0.024 species per five-year period. Yet, from 2000 onwards fewer new species were detected than might be expected given the large increase in human activity across the region in recent decades (IAATO, 2024) and warming climates that likely improve establishment likelihood (Duffy et al., 2017). Indeed, the absolute number of new species detections remains low. These findings suggest that, while new alien species continue to be introduced (Hughes et al., 2015, 2020), the suite of biosecurity policies and practices that have been adopted across the broader Antarctic region over the last three decades have been effective at limiting biological invasions. These policies and practices have included declaration of many of the SOIs as IUCN Category 1 Nature Reserves or their equivalent in domestic legislation, the declaration of many of the islands as World Heritage areas, and strict practices with respect to expectations of the biosecurity procedures of all visitors intending to land on the islands (Table S2; IPBES, 2023). For the Antarctic continent and islands south of 60°S, following the entry into force of the Environmental Protocol in 1998, a consistent focus of the CEP has been on non-native species, in its terminology, and on biosecurity practices (CEP, 2023), but the outcomes are yet to be clearly discerned.

Indeed, regional differences in introduction trends reflect the different levels of human activity and management arrangements across the Antarctic. Across most of the SOIs, models show stable introduction trends, with fewer new aliens observed than expected from 2000 onwards (generally less than 10 species per five year period per subregion). On these islands, human activity is limited by the capacity of scientific stations and research vessels (number of beds), research funding availability, and some limitations on the number of tourist and other landings per annum (De Villiers et al., 2006). Exceptions to these stabilising patterns include the Tristan da Cunha group, Crozet, and Amsterdam and St. Paul Islands, where sampling model expectations of introduction trends continue to increase. Increasing introduction rates along with increasing survey effort trends (Figure 2) suggest that managers in these subregions need to maintain surveillance for new introductions, as well as increase prevention efforts to reduce alien introductions (McGeoch et al., 2023). For SOI subregions with stable alien species trends over time (Figure 2), however, these findings support increasing confidence in the estimated invasion trend (McGeoch et al., 2023).

Differences in long-term average introduction rates across SOI subregions also show, albeit indirectly, the effectiveness of biosecurity practices. Lower average rates of introduction and lower overall numbers of alien species were observed in subregions that were earlier adopters of strict biosecurity procedures, such as the Heard and McDonald Islands and Prince Edward Islands (Table S8). This contrasts with subregions that until recently had largely hortatory biosecurity measures, such as the French Austral Territories (De Villiers et al., 2006), or places with permanent human settlements, such as the Tristan da Cunha group (Table S8).

On the Antarctic continent and its offshore islands (south of 60°S), that are regulated under the Environmental Protocol to the Antarctic Treaty, the situation is less positive. While the continent’s introduction rate has not kept pace with the rapidly increasing number of visitors seen over the last three decades (>100,000 tourist visitors in the 2023/24 season; IAATO, 2024), suggesting some biosecurity success, models show that the introduction rate of alien species continues to increase (Figure 2a). Failure to reverse the introduction trend on the continent may reflect the disparate way that Antarctica is managed, with many countries and private operators working in relative isolation, and with minimal international biosecurity enforcement. Antarctic biosecurity has been criticised for a variety of reasons, including that there is no requirement for alien species monitoring and reporting (Frenot et al., 2005; Hughes et al., 2015; Hughes & Pertierra, 2016; McGeoch et al., 2015), and that there is ongoing non-compliance with existing policies (such as ballast water exchange regulations) and CEP guidelines (Hughes et al., 2020). While there has been some progress (for example, including alien species in Antarctic Specially Protected Areas management plans; Hughes et al., 2021), most of these limitations remain. Since the Protocol came into force in 1998, 40 new alien species (largely inadvertent introductions) have been detected within the Antarctic Treaty area. The north-west Antarctic Peninsula and its offshore islands stand out with most new arrivals on the continent over this period (38 records) (Hughes et al., 2015; Leihy et al., 2023). This is likely a consequence of the highest rates of climate change and global connectivity in the Peninsula region (McCarthy et al., 2022; Siegert et al., 2019), including growing visitor numbers (Hughes et al., 2020).

More generally, several mechanisms could be behind the low or stable rates of introduction across the broader Antarctic region including: few propagules available to become entrained in pre-Antarctic invasion pathways; the Antarctic climate remains too unfavourable for the majority of arriving propagules to establish; or that biosecurity across the region, including in pre-departure stages, has become increasingly effective. Routine biosecurity interceptions of entrained propagules on vessels, aircraft and at Antarctic stations indicate that a broad suite of species is capable of entrainment at pre-Antarctic departure sites (e.g., Houghton et al., 2016). Indeed, despite its perceived isolation, Antarctica is connected to 15% of worldwide ports in the global shipping network (McCarthy et al., 2022). Similarly, Antarctica’s climate barriers in several coastal areas, notably the Antarctic Peninsula, likely provide little current protection from invasion, and are expected to become even less restrictive under future climate change scenarios (Chown et al., 2012; Convey & Peck, 2019; Cuba-Diaz et al., 2023; Duffy et al., 2017). Habitat suitability modelling shows that sites along the Antarctic Peninsula, and extensively across the SOIs, are currently suitable for many of the most impactful globally invasive species and other cold-tolerant species (Duffy et al., 2017).

Against the backdrop of globally increasing rates of alien species introductions (Bonnamour et al., 2021; Roy et al., 2024), and ameliorating polar climates and increasing human visitation across the Antarctic region (Convey & Lebouvier 2009; Hughes et al., 2020), the low and stable rates of introduction found for most of the SOIs suggest that biosecurity has been largely successful at mitigating a further alien species risk. For the Antarctic continent, this is not the case.

Target 6 of the Kunming-Montreal Global Biodiversity Framework includes reducing the introduction of invasive alien species, and monitoring this target by tracking trends in introduction (CBD, 2022). For the SOIs, which fall within national implementation of the provisions of the Convention on Biological Diversity (CBD) (Vicente et al., 2022), this Target largely appears on track to be met by 2030. However, for the Antarctic continent, these global aspirations are not being fully met. Nor are the aspirations of the Antarctic Treaty Consultative Parties that “No species of living organisms not native to the Antarctic Treaty area shall be introduced onto land or ice shelves, or into water, in the Antarctic Treaty area except in accordance with a permit” (Environmental Protocol Annex II Article 4.1 (Antarctic Treaty Secretariat, 1991).

A signal of biosecurity success does not mean that the threat posed by alien species and invasive alien species has abated. Even in places with static introduction trends, new alien species continue to be detected in introduction pathways (e.g., Houghton et al., 2016; Hughes et al., 2015), and most established aliens on the SOIs are not controlled (Convey & Lebouvier, 2009). Prevention of future introductions, through effective biosecurity practices and surveillance of introduction pathways, anticipating future invasions through risk assessments, and developing the preparedness to act when invasions occur (Bergstrom, 2022; De Villiers et al., 2006; Hughes & Pertierra, 2016), will always be key to ongoing conservation success and requires sustained investment. Because the process of invasion is inherently spatially and temporally dynamic, alien species datasets and analyses need to be continually updated as more information becomes available (McGeoch et al., 2023). An open submission system for new or updated alien species records for the Antarctic and SOIs dataset underlying these analyses has recently been released to maintain current information on biological invasions for the region (Leihy et al., 2024). These datasets, along with other findable and accessible alien species checklists (e.g., Dyer et al., 2017; Pagad et al., 2018), may facilitate future assessments on alien species introduction trends and the effectiveness of biosecurity and alien species management policies and practices. As is the case elsewhere, political will, engagement, and education on the risk that invasive species pose to natural environments are also critical to effectively managing alien species risks under what remains a largely hortatory biosecurity system in Antarctica (Hughes & Convey 2010; Hughes & Pertierra, 2016).

Invasive alien species are globally ubiquitous, with an estimated economic impact of US$18.6 billion per year (Diagne et al., 2021). We have demonstrated that, while Antarctica is not exempt from alien species introductions and impacts, the early and stringent international focus on biosecurity in the region has been effective at slowing the rate of new alien species arrivals, especially on the Southern Ocean Islands. These findings provide substantive support for some of the key messages of the recent global assessment of invasive species and their control – that biosecurity policy can drive awareness and effective prevention, that long-term commitment and resourcing support effective policy implementation, and that monitoring policy and management effectiveness provide the evidence that is essential for justifying and refining biosecurity measures (IPBES 2023). For the Antarctic continent itself, largely hortatory policies are failing to deliver the outcomes sought under the Environmental Protocol. Unless mandatory, practicable measures are implemented, changes forecast for the region mediated by biological invasions (Hughes et al., 2020; Rintoul et al., 2018) are likely to be realised.

## Acknowledgements

This work was supported by Australian Research Council SRIEAS Grant SR200100005 Securing Antarctica’s Environmental Future. Laura M. Phillips and Rebecca Hallas provided comments on an earlier version of this manuscript.

## Data availability

All data needed to evaluate the conclusions in the paper are present in the open-access Antarctic and Southern Ocean Islands introduced and invasive alien species dataset (Leihy et al., 2023), and the Supplementary Materials (Table S4).

## Author contributions

Conceptualization: RIL, MAM, DAC, YB, JB, SLC

Data curation: RIL, DAC, LP

Analysis: RIL, LP, YB

Writing—original draft: RIL, MAM, SLC

Writing—review & editing: RIL, MAM, DAC, YB, JB, SLC.

## Supplementary Materials

**Table S1. Glossary of key terms.**

**Table S2. Key Antarctic conservation policy dates.** Overview of human activity and the implementation of key conservation and biosecurity policies across Antarctica and the Southern Ocean Island subregions between 1900 and 2024. These events are expected to have influenced the rate of alien species introductions to the regions due to a change in the occupation of the islands, or the enforcement of a new invasive alien species policy, management plan or reserve status.

**Table S3. Summary of alien species records across the Antarctic and Southern Ocean Island region.**

**Table S4. Additional alien species data records.** Additional alien species records for Antarctica and the Southern Ocean Islands, added to the published and open-access database (Leihy et al., 2023). Occurrence and first record references indicated in the table footnotes, for each new (v.1) and updated (v. 2) record.

**Table S5. Web of Science search terms for survey effort data.** Terms used to conduct a systematic Web of Science search for ecological and environmental publications based on fieldwork conducted in Antarctica and the Southern Ocean Islands between 1900 and 2023, to estimate survey effort over time. For Southern Ocean subregions, papers were sorted manually to exclude publications without a field research component. The temporal trend of relevant publications versus total publications was used to estimate the Antarctic continent search effort proxy, given the large number of Antarctic papers (>30,000) identified in the Web of Science search.

**Table S6. Sampling models selected using the Akaike information criterion (AIC) to chose the best fitting models.** Sampling models were run with the linear and exponential type, and with and without the growth parameter, in the snc function in the alien R package (Buba 2024).

**Table S7. Naïve introduction (A) and survey effort (B) rates for the period from 1900 to 2015 for Antarctica and Southern Ocean Island subregions.** Naïve introduction models assume perfect detection over time. Models fitted with Poisson Generalised Linear Models, p-values corrected for multiple comparisons; p (adjusted) < 0.05*.

**Table S8. Model evaluation metrics.** Mean square error (MSE), bias, and Akaike Information Criterion (AIC) for naïve, Solow and Costello, and sampling models of invasive alien species introduction rate across ten subregions in the Antarctic between 1900 and 2015. Mean and standard deviation for the predicted number of new alien species per five year periods from the naïve introduction rate models.

## References

Antarctic Treaty Secretariat (1991). Protocol on Environmental Protection to the Antarctic Treaty (Madrid, 4 October 1991). Buenos Aires: Antarctic Treaty Secretariat. https://www.ats.aq/e/protocol.html

Auguie, B. (2017). _gridExtra: Miscellaneous Functions for “Grid” Graphics_. R package, version 2.3. https://CRAN.R-project.org/package=gridExtra

Barbraud, C., Des Monstiers, B., Chaigne, A., Marteau, C., Weimerskirch, H., & Delord, K. (2021). Predation by feral cats threatens great albatrosses. Biological Invasions, 23, 2389–2405.

Bartlett, J. C., Convey, P., Newsham, K. K., & Hayward, S. A. L. (2023). Ecological consequences of a single introduced species to the Antarctic: terrestrial impacts of the invasive midge *Eretmoptera murphyi* on Signy Island. Soil Biology and Biochemistry, 180, 108965.

Bergstrom, D. M. (2022). Maintaining Antarctica’s isolation from non-native species. Trends in Ecology and Evolution, 37, 5–9.

Bonnamour, A., Gippet, J. M. W., & Bertelsmeier, C. (2021). Insect and plant invasions follow two waves of globalisation. Ecology Letters 24, 2418–2426.

Buba, Y. (2024). alien: Estimate invasive and alien species (IAS) introduction rates. R package, version 1.0.1. https://cran.r-project.org/web/packages/alien/index.html

Buba, Y., Kiflawi, M., McGeoch, M. A., & Belmaker, J. (2024). Evaluating models for estimating introduction rates of alien species from discovery records. Global Ecology and Biogeography, 33, e13859.

Butchart, S. H., Walpole, M., Collen, B., Van Strien, A., Scharlemann, J. P., Almond, R. E., Baillie, J. E., Bomhard, B., Brown, C., Bruno, J., & Carpenter, K. E. (2010). Global biodiversity: indicators of recent declines. Science 328, 1164–1168.

Chown, S. L., & Language, K. (1994). Recently established Diptera and Lepidoptera on sub-antarctic Marion Island. African Entomology, 2, 57–60.

Chown, S. L., Bergstrom, D. M., Houghton, M., Kiefer, K., Terauds, A., & Leihy, R. I. (2022). Invasive species impacts on sub-Antarctic Collembola support the Antarctic climate-diversity-invasion hypothesis. Soil Biology and Biochemistry, 166, 108579.

Chown, S. L., Huiskes, A. H. L., Gremmen, N. J. M., Lee, J. E., Terauds, A., Crosbie, K., Frenot, Y., Hughes, K. A., Imura, S., Kiefer, K., Lebouvier, M., Raymond, B., Tsujimoto, M., Ware, C., Van de Vijver, B., & Bergstrom, D. M. (2012). Continent-wide risk assessment for the establishment of nonindigenous species in Antarctica. Proceedings of the National Academy of Sciences, 109, 4938–4943.

Chown, S. L., Hull, B., & Gaston, K. J. (2005). Human impacts, energy availability and invasion across Southern Ocean Islands. Global Ecology and Biogeography, 14, 521–528.

Chown, S. L., Brooks, C. M., Terauds, A., Le Bohec, C., van Klaveren-Impagliazzo, C., Whittington, J. D., Butchart, S. H., Coetzee, B. W., Collen, B., Convey, P., Gaston, K. J., Gilbert, N., Gill, M., Hoft, R., Johnston, S., Kennicutt 2nd, M. C., Kriesell, H. J., Le Maho, Y., Lynch, H. J., Palomares, M., Puig-Marco, R., Stoett, P., & McGeoch, M. A. (2017). Antarctica and the strategic plan for biodiversity. PLoS Biology, 15, e2001656.

Committee for Environmental Protection (CEP) (2023). Report of the Twenty-fifth Meeting of the Committee for Environmental Protection (CEP XXV). Helsinki, Finland, May 28 – 1 June 2023.

Committee for Environmental Protection (CEP) (2019). *Non-Native Species Manual*, Revision 2019). Buenos Aires, Argentina.

Connan, M., Schoombie, S., Schoombie, J., Dilley, B., & Ryan, P. G. (2022). Natural recolonisation of sub-Antarctic Marion Island by Common Diving Petrels *Pelecanoides urinatrix*. Ostrich, 93, 271–279.

Convention on Biological Diversity (CBD) (2022). Indicators for the Kunming – Montreal Global Biodiversity Framework. UNEP-WCMC. https://www.gbf-indicators.org

Convey, P., & Lebouvier, M. (2009). Environmental change and human impacts on terrestrial ecosystems of the sub-Antarctic islands between their discovery and the mid-twentieth century. Papers and proceedings of the Royal Society of Tasmania, 143, 33–44.

Convey, P., & Peck, L. S. (2019). Antarctic environmental change and biological responses. Science Advances, 5, eaaz0888.

Cowan, D. A., Chown, S. L., Convey, P., Tuffin, M., Hughes, K., Pointing, S., & Vincent, W. F. (2011). Non-indigenous microorganisms in the Antarctic: assessing the risks. Trends in Microbiology, 19, 540–548.

Cuba-Diaz, M., Fuentes-Lillo, E., Navarrete-Campos, D., & Chwedorzewska, K. J. (2023). Effects of climate change conditions on the individual response and biotic interactions of the native and non-native plants of Antarctica. Polar Biology, 46, 849–863.

De Villiers, M. S., Cooper, J., Carmichael, N., Glass, J. P., Liddle, G. M., McIvor, E., Micol, T., & Roberts, A. (2006). Conservation management at Southern Ocean Islands: towards the development of best-practice guidelines. Polarforschung, 75, 113–131.

Diagne, C., Leroy, B., Vaissiere, A. C., Gozlan, R. E., Roiz, D., Jaric, I., Salles, J. M., Bradshaw, C. J. A., & Courchamp, F. (2021). High and rising economic costs of biological invasions worldwide. Nature, 592, 571–576.

Duffy, G. A., Coetzee, B. W. T., Latombe, G., Akerman, A. H., McGeoch, M. A., & Chown, S. L. (2017). Barriers to globally invasive species are weakening across the Antarctic. Diversity and Distributions, 23, 982–996.

Dyer, E. E., Redding, D. W., & Blackburn, T. M. (2017). The global avian invasions atlas, a database of alien bird distributions worldwide. Scientific Data, 4, 1–12.

Frenot, Y., Chown, S. L., Whinam, J., Selkirk, P. M., Convey, P., Skotnicki, M., & Bergstrom, D. M. (2005). Biological invasions in the Antarctic: extent, impacts and implications. Biological Reviews, 80, 45–72.

Greve, M., Steyn, C., Mathakutha, R., & Chown, S. L. (2017). Terrestrial invasions on sub-Antarctic Marion and Prince Edward islands. Bothalia-African Biodiversity & Conservation, 47, 1–21.

Headland, R. K. (2012). History of exotic terrestrial mammals in Antarctic regions. Polar Records, 48, 123–144.

Houghton, M., McQuillan, P. B., Bergstrom, D. M., Frost, L., Van Den Hoff, J., & Shaw, J. (2016). Pathways of alien invertebrate transfer to the Antarctic region. Polar Biology, 39, 23–33.

Hughes, K. A., Convey, P., & Turner, J. (2021). Developing resilience to climate change impacts in Antarctica: An evaluation of Antarctic Treaty System protected area policy. Environmental Science & Policy, 124, 12–22.

Hughes, K. A., & Convey, P. (2010). The protection of Antarctic terrestrial ecosystems from inter-and intra-continental transfer of non-indigenous species by human activities: a review of current systems and practices. Global Environmental Change, 20, 96–112.

Hughes, K. A., & Pertierra, L. R. (2016). Evaluation of non-native species policy development and implementation within the Antarctic Treaty area. Biological Conservation, 200, 149–159.

Hughes, K. A., Pertierra, L. R., Molina-Montenegro, M. A., & Convey, P. (2015). Biological invasions in terrestrial Antarctica: what is the current status and can we respond? Biodiversity and Conservation, 24, 1031–1055.

Hughes, K. A., Pescott, O. L., Peyton, J., Adriaens, T., Cottier-Cook, E. J., Key, G., Rabitsch, W., Tricarico, E., Barnes, D. K. A., Baxter, N., Belchier, M., Blake, D., Convey, P., Dawson, W., Frohlich, D., Gardiner, L. M., Gonzalez-Moreno, P., James, R., Malumphy, C., Martin, S., Martinou, A. F., Minchin, D., Monaco, A., Moore, N., Morley, S. A., Ross, K., Shanklin, J., Turvey, K., Vaughan, D., Vaux, A. G. C., Werenkraut, V., Winfield, I. J., & Roy, H. E. (2020). Invasive non-native species likely to threaten biodiversity and ecosystems in the Antarctic Peninsula region. Global Change Biology, 26, 2702–2716.

Intergovernmental Science-Policy Platform on Biodiversity and Ecosystem Services (IPBES) (2023). Supplementary material 6.1. Invasive Alien Species in the Antarctic: Policy and Governance. Thematic Assessment Report on Invasive Alien Species and their Control of the Intergovernmental Science-Policy Platform on Biodiversity and Ecosystem Services (pp. 1–10). Bonn, Germany, 2023: IPBES secretariat. 10.5281/zenodo.7430682)

International Association of Antarctic Tour Operators (IAATO) (2024). Data & Statistics. (IAATO, August 19, 2024). https://iaato.org/information-resources/data-statistics/

Lebouvier, M., Laparie, M., Hulle, M., Marais, A., Cozic, Y., Lalouette, L., Vernon, P., Candresse, T., Frenot, Y., & Renault, D. (2011). The significance of the sub-Antarctic Kerguelen Islands for the assessment of the vulnerability of native communities to climate change, alien insect invasions and plant viruses. Biological Invasions, 13, 1195–1208.

Leihy, R. I., Duffy, G. A., & Chown, S. L. (2018). Species richness and turnover among indigenous and introduced plants and insects of the Southern Ocean Islands. Ecosphere, 9, e02358.

Leihy, R. I., Peake, L., Clarke, D. A., Chown, S. L., & McGeoch, M. A. (2023). Introduced and invasive alien species of Antarctica and the Southern Ocean Islands. Scientific Data 10, 200.

Leihy, R. I., Okpokam, N., Chown, S. L., & McGeoch, M. A. (2024). Data submission system - new records of introduced and invasive alien species of Antarctica and the Southern Ocean Islands. LimeSurvey, 8 April 2024. https://aapt.aadpt.cloud.edu.au/limesurvey/

McCarthy, H., Peck, L. S., & Aldridge, D. C. (2022). Ship traffic connects Antarctica’s fragile coasts to worldwide ecosystems. Proceedings of the National Academy of Sciences, 119, e2110303118.

McCarthy, H., Peck, L. S., Hughes, K. A., & Aldridge, D. C. (2019). Antarctica: The final frontier for marine biological invasions. Global Change Biology, 25, 2221–2241.

McClelland, G. T. W., Altwegg, R., van Aarde, R. J., Ferreira, S., Burger, A. E., & Chown, S. L. (2018). Climate change leads to increasing population density and impacts of a key island invader. Ecological Applications, 28, 212–224.

McGeoch, M. A., Buba, Y., Arlé, E., Belmaker, J., Clarke, D. A., Jetz, W., Li, R., Seebens, H., Essl, F., Groom, Q., & García-Berthou, E. (2023). Invasion trends: An interpretable measure of change is needed to support policy targets. Conservation Letters, 16, e12981.

McGeoch, M. A., Shaw, J. D., Terauds, A., Lee, J. E., & Chown, S. L. (2015). Monitoring biological invasion across the broader Antarctic: A baseline and indicator framework. Global Environmental Change, 32, 108–125.

Pagad, S., Genovesi, P., Carnevali, L., Schigel, D., & McGeoch, M. A. (2018). Introducing the global register of introduced and invasive species. Scientific Data 5, 170202.

Plummer, M. (2023)._rjags: Bayesian Graphical Models using MCMC_. R package, version 4–15, https://CRAN.R-project.org/package=rjags

R Core Team, (2022). R: A language and environment for statistical computing. Vienna, Austria: R Foundation for Statistical Computing. https://www.R-project.org/

Rintoul, S. R., Chown, S. L., DeConto, R. M., England, M. H., Fricker, H. A., Masson-Delmotte, V., Naish, T. R., Siegert, M. J., & Xavier, J. C. (2018). Choosing the future of Antarctica. Nature, 558, 233–241.

Rogan-Finnemore, M. (Ed.) (2008). Non-native species in the Antarctic. New Zealand: Gateway Antarctica.

Roy, H. E., Pauchard, A., Stoett, P. J., Truong, T. R., Meyerson, L. A., Bacher, S., Galil, B. S., Hulme, P. E., Ikeda, T., Kavileveettil, S., McGeoch, M. A., Nuñez, M. A., Ordonez, A., Rahlao, S. J., Schwindt, E., Seebens, H., Sheppard, A. W., Vandvik, V., Aleksanyan, A., Ansong, M., August, T., Blanchard, R., Brugnoli, E., Bukombe, J. K., Bwalya, B., Byun, C., Camacho-Cervantes, M., Cassey, P., Castillo, M. L., Courchamp, F., Dehnen-Schmutz, K., Dudeque Zenni, R., Egawa, C., Essl, F., Fayvush, G., Fernandez, R. D., Fernandez, M., Foxcroft, L. C., Genovesi, P., Groom, Q. J., Isabel González, A., Helm, A., Herrera, I., Hiremath, A. J., Howard, P. L., Hui, C., Ikegami, M., Keskin, E., Koyama, A., Ksenofontov, S., Lenzner, B., Lipinskaya, T., Lockwood, J. L., Mangwa, D. C., Martinou, A. F., McDermott, S. M., Morales, C. L., Müllerová, J., Avinash Mungi, N., Munishi, L. K., Ojaveer, H., Pagad, S. N., Pallewatta, N. P. K. T. S., Peacock, L. R., Per, E., Pergl, J., Preda, C., Pyšek, P., Rai, R. K., Ricciardi, A., Richardson, D. M., Riley, S., Rono, B. J., Ryan-Colton, E., Saeedi, H., Shrestha, B. B., Simberloff, D., Tawake, A., Tricarico, E., Vanderhoeven, S., Vicente, J., Vilà, M., Wanzala, W., Werenkraut, V., Weyl, O. L. F., Wilson, J. R. U., Xavier, R. O., & Ziller, S. R. (2024). Curbing the major and growing threats from invasive alien species is urgent and achievable. Nature Ecology and Evolution, 8, 1216–1223.

Russell, J. C., Horn, S. R., Miskelly, C. M., Sagar, R. L., & Taylor, R. H. (2020). Introduced land mammals and their impacts on the birds of the subantarctic Auckland Islands. Notornis, 67, 247–268.

Seebens, H., Blackburn, T. M., Dyer, E. E., Genovesi, P., Hulme, P. E., Jeschke, J. M., Pagad, S., Pyšek, P., Winter, M., Arianoutsou, M., Bacher, S., Blasius, B., Brundu, G., Capinha, C., Celesti-Grapow, L., Dawson, W., Dullinger, S., Fuentes, N., Jäger, H., Kartesz, J., Kenis, M., Kreft, H., Kühn, I., Lenzner, B., Liebhold, A., Mosena, A., Moser, D., Nishino, M., Pearman, D., Pergl, J., Rabitsch, W., Rojas-Sandoval, J., Roques, A., Rorke, S., Rossinelli, S., Roy, H. E., Scalera, R., Schindler, S., Štajerová, K., Tokarska-Guzik, B., van Kleunen, M., Walker, K., Weigelt, P., Yamanaka, T., & Essl, F. (2017). No saturation in the accumulation of alien species worldwide. Nature Communications, 8, 1–9.

Siegert, M., Atkinson, A., Banwell, A., Brandon, M., Convey, P., Davies, B., Downie, R., Edwards, T., Hubbard, B., Marshall, G., Rogelj, J., Rumble, J., Stroeve, J. & Vaughan, D. (2019). The Antarctic Peninsula Under a 1.5°C Global Warming Scenario. Frontiers in Environmental Science, 7, 102.

Solow, A. R., & Costello, C. J. (2004). Estimating the rate of species introductions from the discovery record. Ecology 85, 1822–1825.

Vicente, J. R., Vaz, A. S., Roige, M., Winter, M., Lenzner, B., Clarke, D. A., & McGeoch, M. A. (2022). Existing indicators do not adequately monitor progress toward meeting invasive alien species targets. Conservation Letters, 15, e12918.

Wickham, H. (2016). ggplot2: Elegant Graphics for Data Analysis. New York: Springer-Verlag.

Wonham, M. J., & Pachepsky, E. (2006). A null model of temporal trends in biological invasion records. Ecology Letters, 9, 663–672.

