## Supplementary Tables 1 to 8 for "Antarctic biosecurity policy effectively manages the rates of alien introductions"

**Table S1. Glossary of key terms**

| Term | Synonyms | Definition | Citation |
|---|---|---|---|
| Alien species | adventive, non-indigenous, introduced, exotic, non-native, invasive alien species | An inclusive term for a taxon that has been introduced (either intentionally or accidentally) into an Antarctic area outside of its native range, including synanthropic species that are only associated with sites influenced by humans (e.g., within research stations in Antarctica), OR, a taxon that has spread to a new area beyond its native range from an adjacent introduced population (secondary introductions), including native Antarctic species that have been introduced to new areas within the region by humans, OR, a cryptogenic species of unknown biogeographic origin, in other words species that cannot be ascribed as native or introduced at a particular Antarctic locality. This includes all taxa introduced to places outside their native ranges, including vagrants, domestic species, established or naturalised aliens, invasive alien species, and species that were introduced to areas but have since become locally extinct because they failed to establish a self-sustaining population, died out following establishment, or were eradicated. | Leihy et al., 2023 |
| Detection probability | | Probability of detecting a new alien species at a given time | Buba et al. 2024 |
| Discovery probability | | Probability that the species is first discovered at a given time and not before | |
| Discovery record | | The yearly or cumulative number of discovered alien species (first observation records) over time in a habitat, country, or region | |
| First introduction record | first record | First record of an alien species (or other taxonomic level) discovery at a locality. Here, we used first observation record if available, or estimated first observation record from the publication lag trend between first observation and first publication records from records with both dates available. First introduction records were pooled per subregion so that the earliest introduction date was selected for each alien species within each subregion. | Leihy et al., 2023 |
| First published record | | Year that a new alien species (or other taxonomic level) record was first published in the scientific or grey literature (e.g., in research papers, management documents and reports). This date may involve a publication lag where field observations were not published | |
| First observation record | first observation; discovery records; discovered alien species | Date that an alien species (or other taxonomic level) was first observed (i.e. discovered) at a locality. Often reported as a year or range of years (e.g., "1950- 1953"). May not account for detection or survey effort. | |
| Introduced species data | records | Records of the identity, localities, and dates of introduction of alien species (or other taxonomic levels, such as sub-species or genera). | |

| | | | |
|---|---|---|---|
| Introduction trend | invasion trend | A time series showing the change in the number of alien species in an ecosystem, country, region, or globally | McGeoch et al., 2023 |
| Introduction rate | establishment rate | The rate at which new species are introduced over a particular time period and for a particular region (subnational to global), calculated from the introduction trend. May be called establishment rate if only established species records considered (i.e., excluding vagrants and introduced species that failed to establish self-sustaining populations). | McGeoch et al., 2023 |
| Invasive alien species | | A subset of alien species with an invasive status. That is, an alien species present at a locality where an authoritative sources has described an environmental impact and/or determined that the taxon is widespread, spreading rapid, or present in high abundance. | Pagad et al., 2018 |
| Naïve model | | A model that estimates temporal trends of introduction from the discovery record without accounting for sampling effects. Thus, it implicitly assumes perfect detection of new alien species over time. | Buba et al., 2024 |
| Observed introduction rate | | The rate at which new species are observed (discovered) over a particular time period and for a particular region (subnational to global), calculated from the discovery record (often without accounting for sampling effects). | |
| Publication lag | | Time between when a new alien species is discovered at a locality (i.e., a first observation record) and when the record is published in the scientific or grey literature (i.e., first publication record). | |
| Region | | Here, region refers to the broader Antarctic region, including the Antarctic continent and maritime Antarctic islands (areas south of 60°S, which fall into the Antarctic Treaty area), the sub-Antarctic islands which straddle the Antarctic Polar Front, and several archipelagos with biogeographic and phylogenetic connections to the sub-Antarctic islands in the Southern Ocean (collectively referred to as “Southern Ocean Islands”). Here, the Antarctic region is divided into ten subregions for analysis. | Leihy et al., 2023 |
| Sampling effects | Imperfect detection | Temporal variability in the detection of new alien species due to variability in survey effort | Buba et al., 2024 |
| Sampling model | | A model that accounts for variability in detection probabilities in the estimation of temporal trends of introduction, thereby accounting for sampling effects | |
| Southern Ocean Islands | SOIs | Islands and archipelagos in the Southern Ocean which straggle the Antarctic Polar Front (true sub-Antarctic islands) or have biogeographic and/or phylogenetic affiliations among terrestrial species in the sub-Antarctic. | Leihy et al., 2023 |

| | | | |
|---|---|---|---|
| Solow and Costello model | | A model that estimates temporal trends of introduction from the discovery record, first described by Solow and Costello (2004). It assumes that the overall detection probability increases exponentially over time, and that each species detection also increases due to growth in the population abundance of the species and therefore an increasing detection probability. | Solow & Costello 2004 |
| Subregion | locality | Area or locality for which introduction data is pooled. Here, ten areas or archipelagos within the broader Antarctic region: the Antarctic continent and maritime islands, Île Amsterdam and Île Saint-Paul, Crozet Islands, Heard and McDonald Islands, Kerguelen Islands, Macquarie Island, New Zealand Sub-Antarctic Islands (including Auckland Islands, Campbell Island/Motu Ihupuku, Bounty Islands, The Snares/Tini Heke and the Antipodes Islands), Prince Edward Islands, South Georgia and the South Sandwich Islands, and Tristan da Cunha group (including Gough Islands). | Leihy et al., 2023 |
| Survey effort | Sampling effort, observation effort, discovery process | Investment in surveillance and monitoring activities to observe (and document/report) newly introduced and established species. Here, we use the number of environmental publications with a field research component per year as a proxy for survey effort. This assumes that researchers would report new “unusual” species during the collection of general environmental data, regardless of the scientific discipline. | McGeoch et al., 2023 |

**Table S2. Key Antarctic conservation policy dates.** Overview of human activity and the implementation of key conservation and biosecurity policies across Antarctica and the Southern Ocean Island subregions between 1900 and 2024. These events are expected to have influenced the rate of alien species introductions to the regions due to a change in the occupation of the islands, or the enforcement of a new invasive alien species policy, management plan or reserve status.

| No | Subregion | Year | Event | Citation |
|---|---|---|---|---|
| 1 | Antarctica | 1964 | Agreed Measures for the Conservation of Antarctic Fauna and Flora adopted | ATCM 1964 |
| | | 1982 | Agreed Measures for the Conservation of Antarctic Fauna and Flora entered into force | |
| | | 1991 | Protocol on Environmental Protection to the Antarctic Treaty adopted | Antarctic Treaty Secretariat 1991 |
| | | 1998 | Protocol on Environmental Protection to the Antarctic Treaty entered into force | |
| 2 | Amsterdam & St. Paul | 1949/50 | Martin-de-Vivies station established | Lebourvier & Frenot 2007 |
| | | 1985 | Specific access regulations became effective | |
| | | 2011 | First Management Plan | |
| | | 2017 | National Nature Reserve declared/Second Management Plan | TAAF 2017; Ministère de l'écologie et du développement durable 2022 |
| | | 2019 | World Heritage Listed | World Heritage Committee 2024 |
| 3 | Crozet | 1963/64 | Alfred Faure station established, Ile de la Possession | Lebourvier & Frenot 2007 |
| | | 1985 | Specific access regulations became effective | |
| | | 2006 | National Nature Reserve declared | Ministère de l'écologie et du développement durable 2006 |
| | | 2011 | First Management Plan | |
| | | 2017 | Second Management Plan | TAAF 2017 |
| | | 2019 | World Heritage Listed | World Heritage Committee 2024 |
| 4 | Kerguelen | *1930* | *Whaling ended* | |
| | | 1950/51 | Port-aux-Français station established | Lebourvier & Frenot 2007 |
| | | 1985 | Specific access regulations became effective | |
| | | 2006 | National Nature Reserve declared | Ministère de l'écologie et du développement durable 2006 |
| | | 2011 | First Management Plan | |
| | | 2017 | Management Plan | TAAF 2017 |

| | | | | |
|---|---|---|---|---|
| | | 2019 | World Heritage Listed | World Heritage Committee 2024 |
| 5 | Heard and McDonald | 1947 | ANARE station at Atlas Cove established | World Heritage Committee 2024 |
| | | 1977 | Commonwealth Reserve | |
| | | 1996/97 | First Management Plan/ World Heritage listed | Commonwealth of Australia 2005 |
| | | 2005 | Second Management Plan | |
| 6 | Macquarie | 1933 | Wildlife Sanctuary declared (end of sealing and oil gathering) | |
| | | 1948 | Continuous ANARE visits commenced | Department of Parks, Wildlife and Heritage, Tasmania, 1991; Tasmanian Parks and Wildlife Service 2006 |
| | | 1977/78 | Recognised as a Biosphere Reserve in 1977, and a Nature Reserve 1978, declared a restricted area in 1979 | |
| | | 1991 | First Management Plan | |
| | | 1997 | World Heritage Status | World Heritage Committee 2024 |
| | | 2006 | Second Management Plan | |
| 7 | New Zealand sub-Antarctic islands | 1934 | Auckland Islands declared a Reserve | New Zealand Department of Conservation/Te Papa Atawhai 1997 |
| | | 1953 | Campbell Island declared a Reserve | |
| | | 1961 | Remaining sub-Antarctic islands declared a Reserve | |
| | | 1998 | World Heritage listed | World Heritage Committee 2024 |
| 8 | Prince Edward Islands | 1947 | Continuous South African visits commenced | South African Department of Environmental Affairs and Tourism 1996 |
| | | 1995/96 | Special Nature Reserve Established/ First Management Plan | |
| | | 2010 | Second Management Plan | Centre of Excellence for Invasion Biology 2010 |
| 9 | South Georgia and the South Sandwich Islands | 1965/67 | Whaling industry closed/ Continuous science programs commenced | |
| | | 1975 | Falkland Islands Dependencies Conservation Ordinance | Falkland Islands Dependencies 1975 |
| | | 2000 | First Management Plan | McIntosh & Walton 2000 |
| | | 2011 | Wildlife and Protected Areas Ordinance | GSGSSI 2011 |
| | | 2019 | Biosecurity Handbook 2019-2020 | GSGSSI 2019 |
| 10 | Tristan da Cunha group | 1949/50 | Establishment of commercial fishery and outside administration | Wace & Holdgate 1976 |
| | | 1961 | Settlement evacuated | |
| | | 1963 | Return of settlement 2 years after most evacuated following the volcanic eruption in 1961 near the Tristan Settlement | |

| | | |
|---|---|---|
| 1995/97 | Gough Island World Heritage listed/ Inaccessible Island proclaimed a Nature Reserve | World Heritage Committee 2024 |

**Table S3. Summary of alien species records across the Antarctic and Southern Ocean Island region.**

| No | Subregion | Governance | Description | Number of unique alien records | | Alien species observation period | % alien records missing dates |
|---|---|---|---|---|---|---|---|
| | | | | Total | 1900-2015 | | |
| 1 | Antarctica | Antarctic Treaty System | Continental and maritime Antarctic areas, including all terrestrial and in-land water bodies south of 60°S | 139 | 125 | 1820-2020 | 8.63 |
| 2 | South Georgia and the South Sandwich Islands | United Kingdom (British Overseas territories) | South Georgia and the South Sandwich Islands | 154 | 91 | 1800-2009 | 37.66 |
| 3 | Tristan da Cunha group | | All islands in the archipelago, including Tristan da Cunha, Inaccessible, Nightingale, Gough, Stoltenhoff, and Middle Islands | 373 | 295 | 1793-2010 | 11.80 |
| 4 | Prince Edward Islands | South Africa | All islands in the archipelago, including Marion and Prince Edward Island | 79 | 67 | 1804-2004 | 8.86 |
| 5 | Crozet Islands | France (Terres australes et antarctiques françaises) | All islands in the archipelago, including Île de la Possession, Île de l'Est, Île aux Cochons; Île des Pingouins; Îlots des Apôtres | 172 | 152 | 1821 - 2018 | 7.56 |
| 6 | Kerguelen Islands | | All ~300 islands in the Kerguelen archipelago | 174 | 139 | 1825-2017 | 10.34 |
| 7 | Amsterdam and St. Paul Islands | | Île Saint-Paul and Île d'Amsterdam | 207 | 151 | 1799-2008 | 14.49 |
| 8 | Heard and McDonald Islands | Australia | Heard Island and the McDonald Islands | 7 | 4 | 1855-1992 | 28.57 |
| 9 | Macquarie Island | | Macquarie Island | 103 | 45 | 1810-2014 | 45.63 |

| | | | | | | | |
|---|---|---|---|---|---|---|---|
| 10 | New Zealand sub-Antarctic Islands | New Zealand | New Zealand sub-Antarctic islands, including Campbell/Motu Ihupuku, Auckland islands, Antipode Islands, and The Snares/Tini Heke | 211 | 118 | 1807-1995 | 21.80 |

**Table S4. Additional alien species data records.** Additional alien species records for Antarctica and the Southern Ocean Islands, added to the published and open-access database (Leihy et al., 2023). Occurrence and first record references indicated in the table footnotes, for each new (v.1) and updated (v. 2) record.

| ID | Accepted Name Usage | Verbatim Identification | kingdom | locality | Location Remarks | Occurrence Reference | First Observation Record | First Published Record | First Record Reference | Record Date | Record Note |
|---|---|---|---|---|---|---|---|---|---|---|---|
| 1 | Narcissus L. | *Narcissus sp.* | Plantae | North-west Antarctic Peninsula | East Base, Stonington Island | [385] | 1946 | 1947 | [385] | 9-Oct-23 | v. 1 |
| 2 | Hyacinthoides Heist. ex Fabr. | *Hyacinthoides sp.* | Plantae | North-west Antarctic Peninsula | East Base, Stonington Island | [385] | 1946 | 1947 | [385] | 9-Oct-23 | v. 1 |
| 3 | Lactuca sativa L. | *Lactuca sativa* | Plantae | North-west Antarctic Peninsula | East Base, Stonington Island | [385] | 1946 | 1947 | [385] | 9-Oct-23 | v. 1 |
| 4 | Raphanus raphanistrum L. | *Raphanus raphanistrum* | Plantae | North-west Antarctic Peninsula | East Base, Stonington Island | [385] | 1946 | 1947 | [385] | 9-Oct-23 | v. 1 |
| 5 | Viola tricolor L. | *Viola tricolor* | Plantae | North-west Antarctic Peninsula | East Base, Stonington Island | [385] | 1946 | 1947 | [385] | 9-Oct-23 | v. 1 |
| 6 | Malcolmia maritima (L.) W.T.Aiton | *Malcolmia maritima* | Plantae | North-west Antarctic Peninsula | East Base, Stonington Island | [385] | 1946 | 1947 | [385] | 9-Oct-23 | v. 1 |
| 7 | Eobrachychthonius oudemansi Hammen, 1952 | *Eobranchythonius oudemansi* | Animalia | South Georgia and the South Sandwich Islands | South Sandwich Islands | [174] | 1964 | 1966 | [386] | 9-Oct-23 | v. 2 |
| 8 | Phytomyza syngenesiae (Hardy, 1849) | *Chromatomyia syngenesiae* | Animalia | Antipodes | Antipodes Islands | [9][55] | 1969 | 1976 | [188] | 9-Oct-23 | v. 2 |
| 9 | Phytomyza syngenesiae (Hardy, 1849) | *Chromatomyia syngenesiae* | Animalia | Auckland | Auckland Islands | [9][55] | 1963 | | [188] | 9-Oct-23 | v. 2 |
| 10 | Allium cepa L. | *Allium cepa* | Plantae | New Amsterdam and St. Paul | Île Amsterdam | [340] | | 1986 | [387] | 9-Oct-23 | v. 2 |
| 11 | Allium L. | *Allium porrum* | Plantae | New Amsterdam and St. Paul | Île Amsterdam | [340] | | 1986 | [387] | 9-Oct-23 | v. 2 |

| | | | | | | | | | | | |
|---|---|---|---|---|---|---|---|---|---|---|---|
| 12 | Gladiolus Tourn. ex L. | *Gladiolus sp.* | Plantae | New Amsterdam and St. Paul | Île Amsterdam | [340] | | 1986 | [387] | 9-Oct-23 | v. 2 |
| 13 | Juncus bufonius L. | *Juncus bufonius* | Plantae | New Amsterdam and St. Paul | Île Amsterdam | [65][75][130][340] | 1963 | | [75][130][387] | 9-Oct-23 | v. 2 |
| 14 | Dactylis glomerata L. | *Dactylis glomerata* | Plantae | New Amsterdam and St. Paul | Île Amsterdam | [65][75][130][340] | 1985 | 1986 | [75][130][387] | 9-Oct-23 | v. 2 |
| 15 | Holcus lanatus L. | *Holcus lanatus* | Plantae | New Amsterdam and St. Paul | Île Saint-Paul | [65][75][205][340] | 1857 | 1875 | [75][205][387] | 9-Oct-23 | v. 2 |
| 16 | Lolium perenne L. | *Lolium perenne* | Plantae | New Amsterdam and St. Paul | Île Amsterdam | [65][75][130][340] | 1985 | 1986 | [75][130][387] | 9-Oct-23 | v. 2 |
| 17 | Phragmites australis (Cav.) Trin. ex Steud. | *Phragmites australis* | Plantae | New Amsterdam and St. Paul | Île Amsterdam | [340] | | 1986 | [387] | 9-Oct-23 | v. 2 |
| 18 | Triticum aestivum L. | *Triticum aestivum* | Plantae | New Amsterdam and St. Paul | Île Amsterdam | [130][340] | 1985 | 1986 | [130][387] | 9-Oct-23 | v. 2 |
| 19 | Vulpia bromoides (L.) Gray | *Vulpia bromoides* | Plantae | New Amsterdam and St. Paul | Île Amsterdam | [65][75][130][340] | 1969 | | [75][130][387] | 9-Oct-23 | v. 2 |
| 20 | Canna L. | *Canna sp.* | Plantae | New Amsterdam and St. Paul | Île Amsterdam | [340] | | 1986 | [387] | 9-Oct-23 | v. 2 |
| 21 | Anthriscus cerefolium (L.) Hoffm. | *Anthriscus cerefolium* | Plantae | New Amsterdam and St. Paul | Île Amsterdam | [340] | | 1986 | [387] | 9-Oct-23 | v. 2 |
| 22 | Apiaceae | *Petroselinum sativum* | Plantae | New Amsterdam and St. Paul | Île Amsterdam | [340] | | 1986 | [387] | 9-Oct-23 | v. 2 |
| 23 | Apiaceae | *Petroselinum sativum* | Plantae | New Amsterdam and St. Paul | Île Saint-Paul | [340] | | 1986 | [387] | 9-Oct-23 | v. 2 |
| 24 | Conium maculatum L. | *Conium maculatum* | Plantae | New Amsterdam and St. Paul | Île Amsterdam | [65][75][130][340] | 1985 | 1986 | [75][130][387] | 9-Oct-23 | v. 2 |
| 25 | Daucus carota L. | *Daucus carota* | Plantae | New Amsterdam and St. Paul | Île Amsterdam | [130][340] | 1870 | | [387] | 9-Oct-23 | v. 2 |
| 26 | Foeniculum vulgare Mill. | *Foeniculum dulce* | Plantae | New Amsterdam and St. Paul | Île Amsterdam | [340] | | 1986 | [387] | 9-Oct-23 | v. 2 |
| 27 | Achillea millefolium L. | *Achillea millefolium* | Plantae | New Amsterdam and St. Paul | Île Amsterdam | [130][340] | 1963 | | [130][387] | 9-Oct-23 | v. 2 |

| | | | | | | | | | | | |
|---|---|---|---|---|---|---|---|---|---|---|---|
| 28 | Acmella oleracea (L.) R.K.Jansen | *Spilanthes oleracea* | Plantae | New Amsterdam and St. Paul | Île Amsterdam | [340] | | 1986 | [387] | 9-Oct-23 | v. 2 |
| 29 | Calendula officinalis L. | *Calendula officinalis* | Plantae | New Amsterdam and St. Paul | Île Amsterdam | [340] | | 1986 | [387] | 9-Oct-23 | v. 2 |
| 30 | Cichorium endivia L. | *Cichorium endivia var. latifolia* | Plantae | New Amsterdam and St. Paul | Île Amsterdam | [340] | | 1986 | [387] | 9-Oct-23 | v. 2 |
| 31 | Cichorium intybus L. | *Cichorium intybus* | Plantae | New Amsterdam and St. Paul | Île Amsterdam | [340] | 1960 | | [387] | 9-Oct-23 | v. 2 |
| 32 | Crepis capillaris (L.) Wallr. | *Crepis capillaris* | Plantae | New Amsterdam and St. Paul | Île Amsterdam | [65][130][340] | 1985 | 1986 | [130][387] | 9-Oct-23 | v. 2 |
| 33 | Dahlia variabilis Hort. | *Dahlia variabilis* | Plantae | New Amsterdam and St. Paul | Île Amsterdam | [340] | | 1986 | [387] | 9-Oct-23 | v. 2 |
| 34 | Erigeron canadensis L. | *Conyza canadensis* | Plantae | New Amsterdam and St. Paul | Île Amsterdam | [65][130][340] | 1985 | 1986 | [130][387] | 9-Oct-23 | v. 2 |
| 35 | Lactuca sativa L. | *Lactuca sativa* | Plantae | New Amsterdam and St. Paul | Île Amsterdam | [340] | | 1986 | [387] | 9-Oct-23 | v. 2 |
| 36 | Leontodon taraxacoides Lacaita, 1918 | *Leontodon taraxacoides* | Plantae | New Amsterdam and St. Paul | Île Amsterdam | [65][130][340] | 1985 | 1986 | [130][387] | 9-Oct-23 | v. 2 |
| 37 | Leucanthemum vulgare Lam. | *Leucanthemum vulgare* | Plantae | New Amsterdam and St. Paul | Île Amsterdam | [65][75][340] | 1963 | | [75][130][387] | 9-Oct-23 | v. 2 |
| 38 | Pilosella aurantiaca (L.) F.W.Schultz & Sch.Bip. | *Hieracium aurantiacum* | Plantae | New Amsterdam and St. Paul | Île Amsterdam | [65][130][340] | 1985 | 1986 | [130][387] | 9-Oct-23 | v. 2 |
| 39 | Scolymus maculatus L. | *Scolymus maculatus* | Plantae | New Amsterdam and St. Paul | Île Amsterdam | [340] | | 1986 | [387] | 9-Oct-23 | v. 2 |
| 40 | Sonchus asper (L.) Hill | *Sonchus asper* | Plantae | New Amsterdam and St. Paul | Île Amsterdam | [65][130][340] | 1857 | | [130][387] | 9-Oct-23 | v. 2 |
| 41 | Taraxacum officinale Weber ex Wiggins | *Taraxacum officinale* | Plantae | New Amsterdam and St. Paul | Île Amsterdam | [65][75][130][340] | 1985 | 1986 | [130][387] | 9-Oct-23 | v. 2 |
| 42 | Barbarea verna (Mill.) Asch. | *Barbarea praecox* | Plantae | New Amsterdam and St. Paul | Île Amsterdam | [340] | | 1986 | [387] | 9-Oct-23 | v. 2 |
| 43 | Brassica napus L. | *Brassica napus* | Plantae | New Amsterdam and St. Paul | Île Amsterdam | [65][130][340] | 1960 | | [130][387] | 9-Oct-23 | v. 2 |

| | | | | | | | | | | | |
|---|---|---|---|---|---|---|---|---|---|---|---|
| 44 | Matthiola incana (L.) W.T.Aiton | *Matthiola annua* | Plantae | New Amsterdam and St. Paul | Île Amsterdam | [340] | | 1986 | [387] | 9-Oct-23 | v. 2 |
| 45 | Nasturtium officinale R.Br. | *Nasturtium officinale* | Plantae | New Amsterdam and St. Paul | Île Amsterdam | [340] | | 1986 | [387] | 9-Oct-23 | v. 2 |
| 46 | Raphanus sativus L. | *Raphanus sativus* | Plantae | New Amsterdam and St. Paul | Île Amsterdam | [340] | | 1986 | [387] | 9-Oct-23 | v. 2 |
| 47 | Tropaeolum majus L. | *Tropaeolum majus* | Plantae | New Amsterdam and St. Paul | Île Amsterdam | [65][130][340] | 1985 | 1986 | [130][387] | 9-Oct-23 | v. 2 |
| 48 | Atriplex halimus L. | *Atriplex halimus* | Plantae | New Amsterdam and St. Paul | Île Amsterdam | [130][340] | 1985 | 1986 | [130][387] | 9-Oct-23 | v. 2 |
| 49 | Dianthus barbatus L. | *Dianthus barbatus* | Plantae | New Amsterdam and St. Paul | Île Amsterdam | [340] | | 1986 | [387] | 9-Oct-23 | v. 2 |
| 50 | Dianthus caryophyllus L. | *Dianthus caryophyllus* | Plantae | New Amsterdam and St. Paul | Île Amsterdam | [340] | | 1986 | [387] | 9-Oct-23 | v. 2 |
| 51 | Stellaria media (L.) Vill. | *Stellaria media* | Plantae | New Amsterdam and St. Paul | Île Amsterdam | [65][130][340] | 1948 | | [130][387] | 9-Oct-23 | v. 2 |
| 52 | Mirabilis jalapa L. | *Mirabilis jalapa* | Plantae | New Amsterdam and St. Paul | Île Amsterdam | [340] | | 1986 | [387] | 9-Oct-23 | v. 2 |
| 53 | Rumex acetosella L. | *Rumex acetosella* | Plantae | New Amsterdam and St. Paul | Île Amsterdam | [65][75] [130][340] | 1963 | | [75][130][387] | 9-Oct-23 | v. 2 |
| 54 | Rumex obtusifolius L. | *Rumex obtusifolius* | Plantae | New Amsterdam and St. Paul | Île Amsterdam | [65][130][340] | 1985 | 1986 | [130][387] | 9-Oct-23 | v. 2 |
| 55 | Tamarix gallica L. | *Tamarix gallica* | Plantae | New Amsterdam and St. Paul | Île Amsterdam | [340] | | 1986 | [387] | 9-Oct-23 | v. 2 |
| 56 | Hydrangea macrophylla (Thunb.) Ser. | *Hydrangea hortensis* | Plantae | New Amsterdam and St. Paul | Île Amsterdam | [340] | | 1986 | [387] | 9-Oct-23 | v. 2 |
| 57 | Begonia L. | *Begonia sp.* | Plantae | New Amsterdam and St. Paul | Île Amsterdam | [340] | | 1986 | [387] | 9-Oct-23 | v. 2 |
| 58 | Citrullus lanatus (Thunb.) Matsum. & Nakai | *Citrullus vulgaris* | Plantae | New Amsterdam and St. Paul | Île Amsterdam | [340] | | 1986 | [387] | 9-Oct-23 | v. 2 |
| 59 | Cucumis sativus L. | *Cucumis sativus* | Plantae | New Amsterdam and St. Paul | Île Amsterdam | [340] | | 1986 | [387] | 9-Oct-23 | v. 2 |

| | | | | | | | | | | | |
|---|---|---|---|---|---|---|---|---|---|---|---|
| 60 | Valerianella locusta (L.) Laterr. | *Valerianella olitoria* | Plantae | New Amsterdam and St. Paul | Île Amsterdam | [340] | | 1986 | [387] | 9-Oct-23 | v. 2 |
| 61 | Acacia dealbata Link | *Acacia dealbata* | Plantae | New Amsterdam and St. Paul | Île Amsterdam | [340] | | 1986 | [387] | 9-Oct-23 | v. 2 |
| 62 | Acacia melanoxylon R.Br. | *Mimosa melanoxylon* | Plantae | New Amsterdam and St. Paul | Île Amsterdam | [340] | | 1986 | [387] | 9-Oct-23 | v. 2 |
| 63 | Lathyrus odoratus L. | *Lathyrus odoratus* | Plantae | New Amsterdam and St. Paul | Île Amsterdam | [340] | | 1986 | [387] | 9-Oct-23 | v. 2 |
| 64 | Lotus corniculatus L. | *Lotus corniculatus* | Plantae | New Amsterdam and St. Paul | Île Amsterdam | [65][130][340] | 1964 | | [130][387] | 9-Oct-23 | v. 2 |
| 65 | Phaseolus vulgaris L. | *Phaseolus vulgaris* | Plantae | New Amsterdam and St. Paul | Île Amsterdam | [340] | | 1986 | [387] | 9-Oct-23 | v. 2 |
| 66 | Trifolium dubium Sibth. | *Trifolium dubium* | Plantae | New Amsterdam and St. Paul | Île Amsterdam | [65][75][130][340] | 1985 | 1986 | [75][130][387] | 9-Oct-23 | v. 2 |
| 67 | Trifolium repens L. | *Trifolium repens* | Plantae | New Amsterdam and St. Paul | Île Amsterdam | [65][75][130][340] | 1985 | 1986 | [75][130][387] | 9-Oct-23 | v. 2 |
| 68 | Alnus Mill. | *Alnus sp.* | Plantae | New Amsterdam and St. Paul | Île Amsterdam | [340] | | 1986 | [387] | 9-Oct-23 | v. 2 |
| 69 | Casuarina L. | *Casuarina sp.* | Plantae | New Amsterdam and St. Paul | Île Amsterdam | [340] | | 1986 | [387] | 9-Oct-23 | v. 2 |
| 70 | Geranium robertianum L. | *Geranium robertianum* | Plantae | New Amsterdam and St. Paul | Île Amsterdam | [130][340] | 1985 | 1986 | [130][387] | 9-Oct-23 | v. 2 |
| 71 | Pelargonium zonale (L.) L'Hér. | *Pelargonium zonale* | Plantae | New Amsterdam and St. Paul | Île Amsterdam | [65][130][340] | 1985 | 1986 | [130][387] | 9-Oct-23 | v. 2 |
| 72 | Mentha pulegium L. | *Mentha pulegium* | Plantae | New Amsterdam and St. Paul | Île Amsterdam | [65][130][340] | 1985 | 1986 | [130][387] | 9-Oct-23 | v. 2 |
| 73 | Ocimum basilicum L. | *Ocimum basilicum* | Plantae | New Amsterdam and St. Paul | Île Amsterdam | [340] | | 1986 | [387] | 9-Oct-23 | v. 2 |
| 74 | Prunella vulgaris L. | *Prunella vulgaris* | Plantae | New Amsterdam and St. Paul | Île Amsterdam | [340] | 1985 | 1986 | [75][387] | 9-Oct-23 | v. 2 |
| 75 | Thymus vulgaris L. | *Thymus vulgaris* | Plantae | New Amsterdam and St. Paul | Île Amsterdam | [340] | | 1986 | [387] | 9-Oct-23 | v. 2 |

| | | | | | | | | | | | |
|---|---|---|---|---|---|---|---|---|---|---|---|
| 76 | Plantago lanceolata L. | *Plantago lanceolata* | Plantae | New Amsterdam and St. Paul | Île Amsterdam | [65][75][130][340] | 1960 | | [75][387] | 9-Oct-23 | v. 2 |
| 77 | Verbena officinalis L. | *Verbena officinalis* | Plantae | New Amsterdam and St. Paul | Île Amsterdam | [65][130][340] | 1948 | | [130][387] | 9-Oct-23 | v. 2 |
| 78 | Laurus nobilis L. | *Laurus nobilis* | Plantae | New Amsterdam and St. Paul | Île Amsterdam | [340] | | 1986 | [387] | 9-Oct-23 | v. 2 |
| 79 | Persea americana Mill. | *Persea gratissima* | Plantae | New Amsterdam and St. Paul | Île Amsterdam | [340] | | 1986 | [387] | 9-Oct-23 | v. 2 |
| 80 | Viola tricolor L. | *Viola tricolor* | Plantae | New Amsterdam and St. Paul | Île Amsterdam | [130][340] | 1985 | 1986 | [130][387] | 9-Oct-23 | v. 2 |
| 81 | Malva sylvestris L. | *Malva sylvestris* | Plantae | New Amsterdam and St. Paul | Île Amsterdam | [65][130][340] | 1985 | 1986 | [130][387] | 9-Oct-23 | v. 2 |
| 82 | Eucalyptus L'Hér. | *Eucalyptus sp.* | Plantae | New Amsterdam and St. Paul | Île Amsterdam | [340] | | 1986 | [387] | 9-Oct-23 | v. 2 |
| 83 | Ranunculus repens L. | *Ranunculus repens* | Plantae | New Amsterdam and St. Paul | Île Amsterdam | [130] | 1963 | | [130][387] | 9-Oct-23 | v. 2 |
| 84 | Ficus carica L. | *Ficus carica* | Plantae | New Amsterdam and St. Paul | Île Amsterdam | [340] | | 1986 | [387] | 9-Oct-23 | v. 2 |
| 85 | Cydonia oblonga Mill. | *Cydonia oblonga* | Plantae | New Amsterdam and St. Paul | Île Amsterdam | [340] | | 1986 | [387] | 9-Oct-23 | v. 2 |
| 86 | Fragaria vesca L. | *Fragaria vesca* | Plantae | New Amsterdam and St. Paul | Île Amsterdam | [340] | | 1986 | [387] | 9-Oct-23 | v. 2 |
| 87 | Mespilus germanica L. | *Mespilus germanica* | Plantae | New Amsterdam and St. Paul | Île Amsterdam | [340] | | 1986 | [387] | 9-Oct-23 | v. 2 |
| 88 | Prunus domestica L. | *Prunus domestica* | Plantae | New Amsterdam and St. Paul | Île Amsterdam | [340] | | 1986 | [387] | 9-Oct-23 | v. 2 |
| 89 | Prunus persica (L.) Stokes | *Prunus persica* | Plantae | New Amsterdam and St. Paul | Île Amsterdam | [340] | | 1986 | [387] | 9-Oct-23 | v. 2 |
| 90 | Pyrus malus Murmann | *Pirus malus* | Plantae | New Amsterdam and St. Paul | Île Amsterdam | [340] | | 1986 | [387] | 9-Oct-23 | v. 2 |
| 91 | Rosa L. | *Rosa sp.* | Plantae | New Amsterdam and St. Paul | Île Amsterdam | [340] | | 1986 | [387] | 9-Oct-23 | v. 2 |

| | | | | | | | | | | | |
|---|---|---|---|---|---|---|---|---|---|---|---|
| 92 | Rubus idaeus L. | *Rubus idaeus* | Plantae | New Amsterdam and St. Paul | Île Amsterdam | [340] | | 1986 | [387] | 9-Oct-23 | v. 2 |
| 93 | Ulmus glabra Huds. | *Ulmus glabra* | Plantae | New Amsterdam and St. Paul | Île Amsterdam | [340] | | 1986 | [387] | 9-Oct-23 | v. 2 |
| 94 | Citrus aurantium L. | *Citrus sinensis* | Plantae | New Amsterdam and St. Paul | Île Amsterdam | [340] | | 1986 | [387] | 9-Oct-23 | v. 2 |
| 95 | Acer pseudoplatanus L. | *Acer pseudoplatanus* | Plantae | New Amsterdam and St. Paul | Île Amsterdam | [340] | | 1986 | [387] | 9-Oct-23 | v. 2 |
| 96 | Lycopersicum esculentum Mill | *Lycopersicum esculentum* | Plantae | New Amsterdam and St. Paul | Île Amsterdam | [340] | | 1986 | [387] | 9-Oct-23 | v. 2 |
| 97 | Vitis x vinifera L. | *Vitis vinifera* | Plantae | New Amsterdam and St. Paul | Île Amsterdam | [340] | | 1986 | [387] | 9-Oct-23 | v. 2 |
| 98 | Cryptomeria japonica (Thunb. ex L.f.) D.Don | *Cryptomeria japonica* | Plantae | New Amsterdam and St. Paul | Île Amsterdam | [340] | | 1986 | [387] | 9-Oct-23 | v. 2 |
| 99 | Cupressus macrocarpa Hartw. ex Gordon | *Cupressus macrocarpa* | Plantae | New Amsterdam and St. Paul | Île Amsterdam | [340] | | 1986 | [387] | 9-Oct-23 | v. 2 |
| 100 | Thuja occidentalis L. | *Thuya occidentalis* | Plantae | New Amsterdam and St. Paul | Île Amsterdam | [340] | | 1986 | [387] | 9-Oct-23 | v. 2 |
| 101 | Pinus halepensis Mill. | *Pinus halepensis* | Plantae | New Amsterdam and St. Paul | Île Amsterdam | [340] | | 1986 | [387] | 9-Oct-23 | v. 2 |
| 102 | Pinus pinaster Aiton | *Pinus pinaster* | Plantae | New Amsterdam and St. Paul | Île Amsterdam | [340] | | 1986 | [387] | 9-Oct-23 | v. 2 |
| 103 | Cartodere nodifer (Westwood, 1839) | *Lathridius nodifer* | Animalia | East Antarctica | Davis Station | [388] | 1973 | 1978 | [388] | 23/10/2023 | v. 2 |
| 104 | Tetragnatha nitens (Audouin, 1826) | *Tetragnatha gulosa* | Animalia | New Amsterdam and St. Paul | Île Saint-Paul | [23][45] | 1899 | 1903 | [394] | 13-Nov-23 | v. 2 |
| 105 | Theridium tepidariorum Koch, 1841 | *Theridium tepidariorum* | Animalia | New Amsterdam and St. Paul | Île Saint-Paul | [23][45][340] | 1857-1859 | 1872 | [394] | 13-Nov-23 | v. 2 |
| 106 | Cryptops megaloporus Haase, 1887 | *Cryptops (Cryptops) megaloporus* | Animalia | Auckland | Auckland Islands | [139] | | 1887 | [389] | 13-Nov-23 | v. 2 |

| | | | | | | | | | | | |
|---|---|---|---|---|---|---|---|---|---|---|---|
| 107 | Henicops maculatus Newport, 1845 | *Henicops maculatus* | Animalia | Campbell | Campbell Island/Motu Ihupuku | [139] | | 1964 | [139] | 13-Nov-23 | v. 2 |
| 108 | Cormocephalus Newport, 1844 | *Cormocephalus sp.* | Animalia | Campbell | Campbell Island/Motu Ihupuku | [139] | | 1909 | [390] | 13-Nov-23 | v. 2 |
| 109 | Lamyctes africanus Porat, 1871 | *Lamyctes (Metalamyctes) africanus* | Animalia | New Amsterdam and St. Paul | Île Saint-Paul | [139][340] | | 1907 | [391] | 13-Nov-23 | v. 2 |
| 110 | Lithobius Leach, 1814 | *Lithobius sp.* | Animalia | New Amsterdam and St. Paul | Île Saint-Paul | [139] | 1938 | 1940 | [392] | 13-Nov-23 | v. 2 |
| 111 | Geophilus flavus De Geer, 1778 | *Necropholcophagus longicornis* | Animalia | Tristan group | Tristan da Cunha Island | [3][139] | | 1928 | [393] | 13-Nov-23 | v. 2 |
| 112 | Lamyctes emarginatus Newport, 1844 | *Lamyctes fulvicornis* | Animalia | Tristan group | Tristan da Cunha Island | [3][139] | | 1956 | [139] | 13-Nov-23 | v. 2 |
| 113 | Lithobius Leach, 1814 | *Lithobius sp.* | Animalia | Tristan group | Tristan da Cunha Island | [139] | | 1956 | [139] | 13-Nov-23 | v. 2 |
| 114 | Opogona omoscopa (Meyrick, 1893) | *Exala strassenella* | Animalia | New Amsterdam and St. Paul | Île Amsterdam | [55][340] | 1899 | 1903 | [197] | 13-Nov-23 | v. 2 |

Data references: [3] Holdgate, M. (1965) Philos. Trans. R. Soc. B.; [9] Marris, J. W. M. (2000) J. R. Soc. N. Z.; [23] Strand, E. (1909) Deutsche Südpolar-Expedition; [45] Berland, L. (1947) Mém. Mus. natl. hist. nat.; [55] Chown, S. L. & Convey, P. (2016) Annu. Rev. Entomol.; [65] Shaw, J. D. (2010); [75] TAAF (2017) Plan de gestion 2018-2027; [130] Frenot, Y., et al. (2001) Biol. Conserv.; [139] Pugh, P. J. A. (2013) J. Nat. Hist.; [174] Pugh, P. J. A. (1994) Zool. J. Linn. Soc.; [188] Harrison, R. A. (1976) J. R. Soc. N. Z.; [197] Viette, P. (1959) Le Bulletin de la SEF; [205] Hooker, J. D. (1875) Bot. J. Linn.; [340] Marchand, D. (1997); [385] Bingham, E. (1947) Pol. Rec.; [386] Wallwork, J. A. (1966) BAS Bulletin; [387] Jolinon, J.C. (1986) C.N.F.R.A.; [388] Rounsevell, D. (1978) Pac. Insects; [389] Haase, E. (1887) Abh. Ber. Mus. Tierk. Volkerk. Dresden.; [390] Benham, W.B. (1909) The Subantarctic islands of New Zealand; [391] Attems, C. (1907) Deutsche Südpolar-Expedition; [392] Attems, C. (1940) Mem. Mus. Paris. NS.; [393] Attems, C. (1928) Ann. S. Afr. Mus.; [394] Enderlein, G. (1903) Wiss. Erg. Deut. Tiefsee-Exp. Valdivia.

**Table S5. Web of Science search terms for survey effort data.** Terms used to conduct a systematic Web of Science search for ecological and environmental publications based on fieldwork conducted in Antarctica and the Southern Ocean Islands between 1900 and 2023, to estimate survey effort over time. For Southern Ocean subregions, papers were sorted manually to exclude publications without a field research component. The temporal trend of relevant publications versus total publications was used to estimate the Antarctic continent search effort proxy, given the large number of Antarctic papers (>30,000) identified in the Web of Science search.

| Subregion | Search terms | N. papers | Search date |
|---|---|---|---|
| Antarctica | ”South Orkney*” OR ”Signy Isl*” OR ”Laurie Isl*” OR ”Coronation Isl*” OR ”Powell Isl*” ”South Shetland*” OR ”Deception Isl*” OR ”King George Isl*” OR ”Fildes Penin*” OR ”Livingston Isl*” OR ”Elephant Isl*” OR “Greenwich Isl*” OR ”Nelson Isl*” OR “Half Moon Isl*” OR “Ardley Isl*” OR “Robert Isl*” ”Antarc* Peninsula” OR ”Palmer Land” OR “Graham Land” OR “Seymour Isl*” OR “Marambio Isl*” OR “Anvers Isl*” OR “Adelaide Isl*” OR “Galindez Isl*” OR “Melchior Isl*” OR “Dundee Isl*” OR “Goudier Isl*” OR “Horseshoe Isl*” OR “Doumer Isl*” ”McMurdo Dry Valley*” OR ”Ross Sea region” ”Vestfold Hills” OR “Bunger Hills” OR “Larsemann Hills” OR “Schirmacher Oasis” ”Transantarctic mountain*” OR ”Ellsworth Land” OR ”Marie Byrd Land” OR “King Edward VII Land” ”Victoria Land” OR ”Oates Land” OR ”George V Land” OR “Ross Isl*” ”Terre Adélie” OR ”Terre Adelie” OR “Wilkes Land” OR ”Queen Mary Land” OR ”Wilhelm II Land” OR ”Princess Elizabeth Land” OR “Adélie Land” ”Macrobertson Land” OR ”Kemp Land” OR ”Dronning Maud Land” OR ”Queen Maud Land” OR ”Enderby Land” OR “Coats Land” OR “Queen Elizabeth Land” OR “Vesleskarvet” OR “East Ongul Isl*” OR "Sør Rondane Mountain*" | 30,759 | 30/8/2023 |
| South Georgia and the South Sandwich islands | TS=”South Georgia” OR TS=”South Georgia Isl*” OR TS=”Bird Isl*”<br>TS=”South Sandwich Isl*” | South Georgia: 2437<br>South Sandwich: 2848 | 18/4/2023 |
| Tristan da Cunha group | TS=”Tristan da Cunha” OR TS=”Nightingale Isl*” OR TS=”Inaccessible Isl*” OR TS=”Gough Isl*” OR TS = “Tristan da Cunha Isl*” OR TS=“Tristan da Cunha Archipel*” | 740 | |
| Prince Edward Islands | TS=”Marion Isl*” OR TS=”Prince Edward Islands” OR TS=(Marion AND Prince Edward) | 1031 | |
| Crozet archipelago | TS=”Crozet Isl*” OR TS=”Iles Crozet” OR TS=”Possession Isl*” OR TS=”Ile aux Cochon” OR TS=”Ile de l’Est” OR TS=”Ile des Pinguoins” or TS=”Ilots | 481 | |

| | | |
|---|---|---|
| | des Apôtres” OR TS=”Crozet Archipel*” OR TS=”Île aux Cochon” OR TS=”Île de l’Est” OR TS=”Île des Pinguoins” or TS=”Îlots des Apôtres” OR TS=” Îles Crozet” | |
| Kerguelen Islands | TS=”Kerguelen Isl*” OR TS=”Kerguelen Archipel*” OR TS=”Iles Kerguelen” OR TS=“Îles Kerguelen” | 1076 |
| Amsterdam and St. Paul | TS=”Amsterdam Isl” OR TS=”St. Paul Isl*” OR TS=”Ile Amsterdam” OR TS= “Île Amsterdam” OR TS=”Île Saint-Paul” OR TS=”Ile Saint-Paul” OR TS=”Saint Paul Isl*” OR TS=”New Amsterdam Isl*” OR TS=”Nouvelle Amsterdam” | 571 |
| Heard and McDonald Islands | TS=”Heard Isl* OR TS=”McDonald Isl*” OR TS=”Heard and McDonald Isl*” | 319 |
| Macquarie Island | TS=”Macquarie Isl*” OR TS=”Bishop and Clerk Isl*” | 2534 |
| New Zealand sub-Antarctic islands | TS=”Campbell Isl*” OR TS=”Auckland Isl*” OR TS=”Antipodes Isl*” OR TS=”Snares Isl*” OR TS=”Bounty Isl*” OR TS=”The Snares” OR TS=”Motu Ihupuku” OR TS=”Tini Heke” | 1890 |

**Table S6. Sampling models selected using the *Akaike information criterion* (*AIC*) to chose the best fitting models.** Sampling models were run with the linear and exponential type, and with and without the growth parameter, in the snc function in the alien R package (Buba, 2024).

| **Subregion** | **Introduction rate model type** | **$\gamma_2$ included** | **AIC** |
|---|---|---|---|
| Antarctica | linear | no | 211.54 |
| Amsterdam & St. Paul | exponential | no | 352.01 |
| Crozet | linear | yes | 353.06 |
| Heard & McDonald islands | *Models failed to converge* | | |
| Kerguelen | exponential | no | 222.82 |
| Macquarie | linear | no | -18.01 |
| New Zealand islands | exponential | no | 280.71 |
| Prince Edward islands | exponential | yes | 38.01 |
| South Georgia & South Sandwich | *Models failed to converge* | | |
| Tristan group | exponential | no | 938.10 |

**Table S7. Naïve introduction (a) and survey effort (b) rates for the period from 1900 to 2015 for Antarctica and Southern Ocean Island subregions.** Naïve introduction models assume perfect detection over time. Models fitted with Poisson Generalised Linear Models, p-values corrected for multiple comparisons; p (adjusted) < 0.05*.

**a) Invasive alien species trends over time (naïve model)**

| Subregion | Estimate ($\beta_1$) | SE | Intercept | z | P |
|---|---|---|---|---|---|
| Antarctica | 0.02 | <0.01 | 0.58 | 5.72 | <0.001* |
| Crozet archipelago | 0.02 | <0.01 | 0.37 | 7.81 | <0.001* |
| Heard and McDonald Islands | 0.02 | 0.02 | -3.39 | 1.35 | 0.885 |
| Kerguelen Islands | <0.01 | <0.01 | 1.58 | 1.39 | 0.824 |
| Macquarie Island | <0.01 | <0.01 | 0.18 | 1.78 | 0.743 |
| Amsterdam and St. Paul | 0.02 | <0.01 | 0.82 | 6.20 | <0.001* |
| New Zealand sub-Antarctic islands | <0.01 | <0.01 | 1.89 | -1.72 | 0.430 |
| Prince Edward Islands | 0.02 | <0.01 | 0.12 | 3.79 | 0.001* |
| South Georgia and the South Sandwich islands | <0.01 | <0.01 | 1.30 | 0.43 | 1.000 |
| Tristan da Cunha group | <0.01 | <0.01 | 2.25 | 2.90 | 0.019* |

**b) Survey effort trends over time**

| Subregion | Estimate | SE | Intercept | z | P |
|---|---|---|---|---|---|
| Antarctica | 0.07 | <0.01 | -0.11 | 86.60 | <0.001* |
| Crozet archipelago | 0.06 | <0.01 | -2.57 | 14.79 | <0.001* |
| Heard and McDonald Islands | 0.04 | <0.01 | -1.65 | 10.12 | <0.001* |
| Kerguelen Islands | 0.06 | <0.01 | -2.48 | 18.27 | <0.001* |
| Macquarie Island | 0.04 | <0.01 | 0.42 | 19.16 | <0.001* |
| Amsterdam and St. Paul | 0.03 | <0.01 | -0.94 | 6.24 | <0.001* |
| New Zealand sub-Antarctic islands | 0.03 | <0.01 | 0.88 | 21.28 | <0.001* |
| Prince Edward Islands | 0.06 | <0.01 | -2.38 | 19.90 | <0.001* |
| South Georgia and the South Sandwich islands | 0.05 | <0.01 | -0.83 | 23.30 | <0.001* |

| | | | | | |
|---|---|---|---|---|---|
| Tristan da Cunha group | 0.05 | <0.01 | -1.68 | 13.31 | <0.001* |

**Table S8. Model evaluation metrics.** Mean square error (MSE), bias, and Akaike Information Criterion (AIC) for naïve, Solow and Costello, and sampling models of invasive alien species introduction rate across ten subregions in the Antarctic between 1900 and 2015. Mean and standard deviation for the predicted number of new alien species per five year periods from the naïve introduction rate models.

| | **Naïve** | | | | **Solow & Costello** | | **Sampling** | | |
|---|---|---|---|---|---|---|---|---|---|
| **Sub-region** | **MSE** | **Bias** | **AIC** | **Mean (SD) number of predicted new aliens per five-years** | **MSE** | **Bias** | **MSE** | **Bias** | **AIC** |
| Antarctica | 61.58 | <0.01 | 237.27 | 5.35 (3.00) | 61.29 | -0.57 | 60.17 | -0.48 | 211.54 |
| Amsterdam & St. Paul | 155.39 | <0.01 | 345.49 | 6.57 (3.59) | 1.93E+04 | -81.18 | 177.01 | -1.58 | 352.01 |
| Crozet | 100.16 | <0.01 | 279.48 | 6.48 (4.69) | 3.00E+04 | -72.09 | 103.77 | -3.60 | 353.06 |
| Heard & McDonald islands | 0.22 | <0.01 | 25.21 | 0.17 (0.13) | 44.32 | -2.90 | NA | NA | NA |
| Kerguelen | 94.20 | <0.01 | 307.93 | 5.96 (0.72) | 2.29E+109 | -1.00E+54 | 97.45 | <0.01 | 222.82 |
| Macquarie | 4.85 | <0.01 | 94.93 | 1.96 (0.54) | 6.84E+08 | -1.10E+04 | 5.17 | <0.01 | -18.01 |
| New Zealand islands | 86.05 | <0.01 | 321.23 | 5.13 (0.83) | 86.05 | <0.01 | 151.23 | <0.01 | 280.71 |
| Prince Edward islands | 16.30 | <0.01 | 146.45 | 2.91 (1.45) | 301.41 | -10.63 | 16.27 | -0.52 | 38.01 |
| South Georgia & South Sandwich | 44.11 | <0.01 | 212.35 | 3.96 (0.18) | 43.59 | -0.05 | NA | NA | NA |
| Tristan group | 271.16 | <0.01 | 489.77 | 12.83 (2.22) | 9.20E+64 | -6.58E+31 | 281.46 | -1.28 | 938.10 |

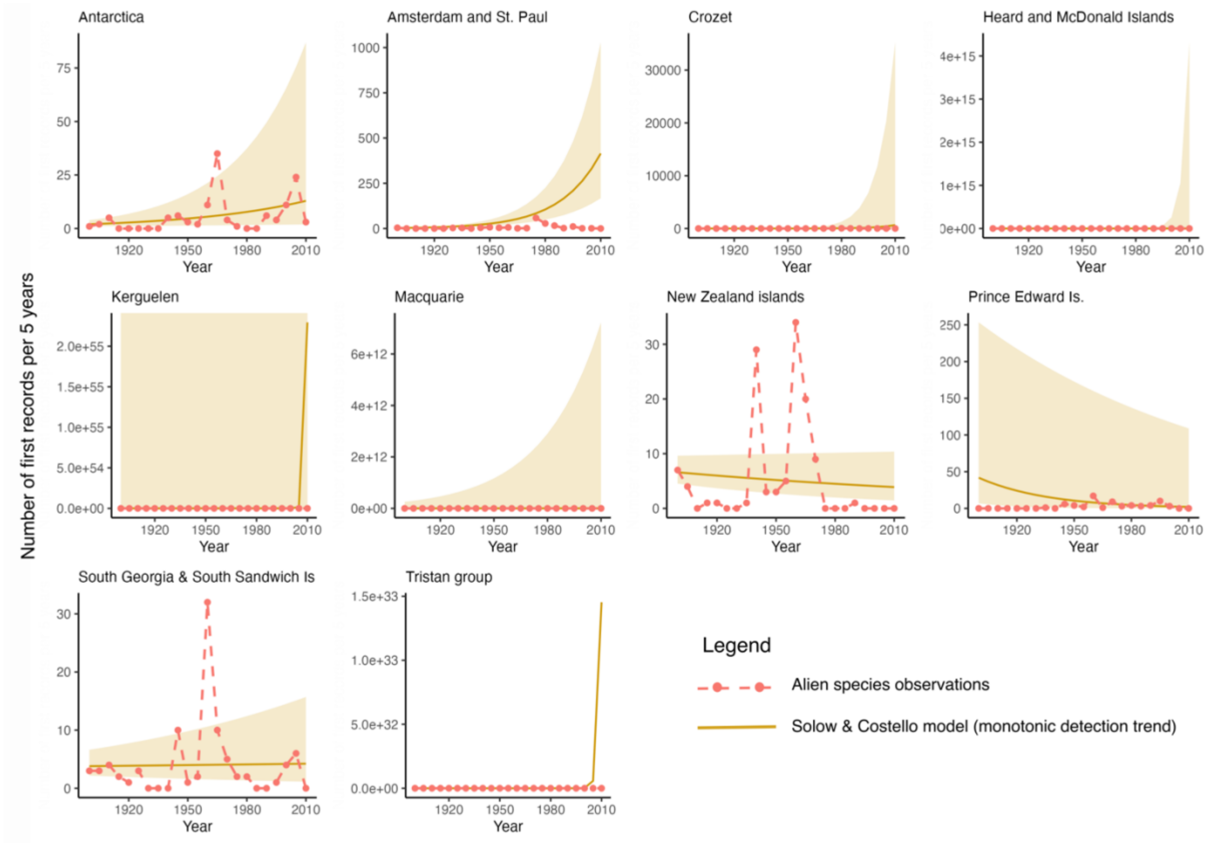


**Figure S1. Solow and Costello model.** Solow and Costello model (yellow lines and shaded 95% confidence intervals) which assumes that survey effort changes monotonically with time and that the underlying introduction rate changes exponentially with time for Antarctic and Southern Ocean Island subregions. Red points and dashed lines indicate the discovery record. For most subregions, the Solow and Costello model was a poor fit to the observed introduction data compared to the naïve model which assumes perfect detection, with larger bias and, in most subregions, very large 95% confidence intervals (Table S4). For the New Zealand sub-Antarctic islands and South Georgia and the South Sandwich Islands, the Solow and Costello model resembles the naïve model.

**Supplementary references**

Antarctic Treaty Consultative Meeting (ATCM) (1964). Agreed Measures for the Conservation of Antarctic Fauna and Flora. Antarctic Treaty Consultative Parties. https://documents.ats.aq/recatt/att080_e.pdf [Accessed 30 October 2023]

Buba, Y. (2024). alien: Estimate invasive and alien species (IAS) introduction rates. R package, version 1.0.1. https://cran.r-project.org/web/packages/alien/index.html

Buba, Y., Kiflawi, M., McGeoch, M. A., & Belmaker, J. (2024). Evaluating models for estimating introduction rates of alien species from discovery records. *Global Ecology and Biogeography*, 33, e13859.

Centre of Excellence for Invasion Biology (CIB) (2010). Prince Edward Islands Environmental Management Plan. Stellenbosch University, Matieland: DST-NRF Centre of Excellence for Invasion Biology, version 0.2., 2010. https://www.environment.gov.za/documents/strategicdocuments [Accessed April 2020]

Commonwealth of Australia (2005). *Heard Island and McDonald Islands Marine Reserve Management Plan*. Commonwealth of Australia.

Department of Parks, Wildlife and Heritage (1991). *Macquarie Island Nature Reserve Management Plan*. Tasmania: Department of Parks, Wildlife and Heritage, Tasmania. https://nla.gov.au/nla.obj-2656572536/view [Accessed 30 October 2023]

Falkland Islands Dependencies (1975). Conservation Ordinance. *Falkland Islands Gazette*, DS 1, 17-21.

Government of South Georgia & the South Sandwich Islands (GSGSSI) (2019). Biosecurity Handbook 2019-2020. Stanley, Falkland Islands: Government of South Georgia & the South Sandwich Islands.

Government of South Georgia & the South Sandwich Islands (GSGSSI) (2013). Wildlife and Protected Areas (Amendment) Ordinance 2013. Stanley, Falkland Islands: Government of South Georgia & the South Sandwich Islands (2011, amended 2013).

Lebourvier, M. & Frenot, Y. (2007). Conservation and Management in the French Sub-Antarctic Islands and surrounding seas. *Papers and proceedings of the Royal Society of Tasmania*, 141, 23-28.

Leihy, R. I., Peake, L., Clarke, D. A., Chown, S. L., & McGeoch, M. A. (2023). Introduced and invasive alien species of Antarctica and the Southern Ocean Islands. *Scientific Data,* 10, 200.

McIntosh, E. & Walton, D. W. H. (2000). *Environmental Management Plan for South Georgia*. Government of South Georgia and the South Sandwich Islands.

Ministère de l'écologie et du développement durable (2022). Décret n° 2022-157 du 10 février 2022 portant extension et modification de la réglementation de la réserve naturelle nationale des Terres australes françaises. https://www.legifrance.gouv.fr/loda/id/LEGIARTI000045156144/2022-02-12#LEGIARTI000045156144 [Accessed 30 October 2023]

Ministère de l'écologie et du développement durable (2006). Décret no 2006-1211 du 3 octobre 2006 portant création de la réserve naturelle des Terres australes françaises, Paris. https://reserve-australes.taaf.fr/gestion-et-reglementation/cadre-reglementaire/ [Accessed April 2020]

New Zealand Department of Conservation/Te Papa Atawhai (1997). *Subantarctic Island Heritage. Nomination of the New Zealand sub-Antarctic Islands by the Government of New Zealand for inclusion in the World Heritage List*. Wellington, New Zealand).

South African Department of Environmental Affairs and Tourism (1996). Prince Edward Islands Management Plan (ISBN 0-621-17584-6).

Tasmanian Parks and Wildlife Service (2006). Macquarie Island Nature Reserve and World Heritage Area Management Plan 2006. Hobart: Parks and Wildlife Service, Department of Tourism, Arts and the Environment.

Terres Australes et Antarctiques Françaises (TAAF) (2017). *Plan de gestion 2018-2027 de la réserve naturelle nationale des Terres australes françaises*. Terres Australes et Antarctiques Françaises.

Wace, N. M., & Holdgate, M. W. (1976). Man and Nature in the Tristan da Cunha Islands. International Union for Conservation of Nature and Natural Resources.

World Heritage Committee (2024). *World Heritage List*. https://whc.unesco.org/en/list/?&mode=table [Accessed online 22 April 2024]
